# ERK basal and pulsatile activity are differentially regulated in mammalian epidermis to control proliferation and exit from the stem cell compartment

**DOI:** 10.1101/2020.04.12.038406

**Authors:** Toru Hiratsuka, Ignacio Bordeu, Gunnar Pruessner, Fiona M. Watt

## Abstract

Fluctuation in signal transduction pathways is frequently observed during mammalian development. However, its role in regulating stem cells has not been explored. Here we tracked spatiotemporal ERK MAPK dynamics in human epidermal stem cells. While stem cells and differentiated cells were distinguished by high and low stable basal ERK activity, respectively, we also found cells with pulsatile ERK activity. Transitions from Basal^hi^-Pulse^lo^ (stem) to Basal^hi^-Pulse^hi^, Basal^mid^-Pulse^hi^, and Basal^lo^-Pulse^lo^ (differentiated) cells occurred in expanding keratinocyte colonies and in response to a range of differentiation stimuli. Pharmacological inhibition of ERK induced differentiation only when cells were in the Basal^mid^-Pulse^hi^ state. Basal ERK activity and pulses were differentially regulated by DUSP10 and DUSP6, leading us to speculate that DUSP6-mediated ERK pulse downregulation promotes initiation of differentiation whereas DUSP10-mediated downregulation of mean ERK activity promotes and stabilizes post-commitment differentiation. Quantification of MAPK1/3, DUSP6 and DUSP10 transcripts in individual cells demonstrated that ERK activity is controlled both transcriptionally and post-transcriptionally. When cells were cultured on a topography that mimics the epidermal-dermal interface, spatial segregation of mean ERK activity and pulses was observed. In vivo imaging of mouse epidermis revealed a patterned distribution of basal cells with pulsatile ERK activity and downregulation was linked to the onset of differentiation. Our findings demonstrate that ERK MAPK signal fluctuations link kinase activity to stem cell dynamics.

**Significance:** Understanding how intracellular signaling cascades control cell fate is a key issue in stem cell biology. Here we show that exit from the stem cell compartment in mammalian epidermis is characterised by pulsatile ERK MAPK activity. Basal activity and pulses are differentially regulated by DUSP10 and DUSP6, two phosphatases that have been shown previously to regulate differentiation commitment in the epidermis. ERK activity is controlled both transcriptionally and post-transcriptionally. Spatial segregation of mean ERK activity and pulses is observed both in reconstituted human epidermis and in mouse epidermis. Our findings demonstrate the tight spatial and temporal regulation of ERK MAPK expression and activity in mammalian epidermis.

## Introduction

Fluctuation in signals involving Notch, Wnt, FGF, p53, NF-κB and other pathways plays a significant role in mammalian development and physiology(1-6). Spatial and temporal heterogeneity of the signalling profiles in individual cells regulates gene expression and cell fate. This heterogeneity reflects changes in the intracellular and extracellular environment, including biochemical noise in signalling components, the availability and gradient of growth factors, and interactions with surrounding cells.

Spatiotemporal activation patterns of Extracellular signal-Regulated Kinase (ERK) MAPK are believed to play a significant role in a variety of cellular processes, including cell proliferation, migration and differentiation(7, 8). Recent single-cell live imaging approaches have revealed fluctuating and propagating features of ERK activation(9-13). In the epidermis of living mice, bursts of ERK activity radially propagate to neighbouring cells and can be triggered by wounding and other external stimuli(11).

Stem cells reside in the basal layer of the epidermis, where they self-renew or generate committed cells that undergo terminal differentiation. Stem cell markers include high levels of α6 and β1 integrins(14, 15), Delta-like 1 (DLL1), and Lrig1(16, 17). Despite the requirement of ERK activity to maintain epidermal stem cells(18-22), its role in cell state transitions, such as proliferative/quiescent and differentiation commitment, have not been studied.

Here we show, by live imaging of thousands of human epidermal cells, that there are dynamic transitions in ERK activity during stem cell colony expansion and differentiation. ERK pulse activity and basal levels are independently regulated by DUSP6 and DUSP10, components of the autoregulatory protein phosphatase network that acts as a switch between the stem cell state and the differentiated cell state(23). We also observe spatial segregation of cells with different ERK temporal patterns on substrates mimicking human dermis and in live mouse skin, establishing the physiological significance of our observations.

## Results

### Transitions in ERK activity dynamics during expansion of stem cell colonies

We live imaged ERK activity in individual primary human neonatal keratinocytes (HNKs) via lentiviral expression of a nuclear-tagged FRET biosensor for ERK, EKAR-EVnls(24). NHKs were seeded on a 3T3 feeder layer(25) and expanding colonies were subjected to live imaging on different days from 3 to 8 days after plating (Fig. 1 *A-C*). Within colonies, mean ERK activity (hereinafter referred to as “basal activity”) was lower in large (median in Fig. 1*C*) than small keratinocytes. Keratinocytes are known to enlarge as they undergo differentiation, and the proportion of differentiated cells increases as colonies expand. Therefore this observation is consistent with the previously reported downregulation of basal ERK activity in differentiated cells(14). We also observed that there was a wider range of ERK activity in smaller cells than large cells (whiskers in Fig. 1*C*).

**Fig. 1.**
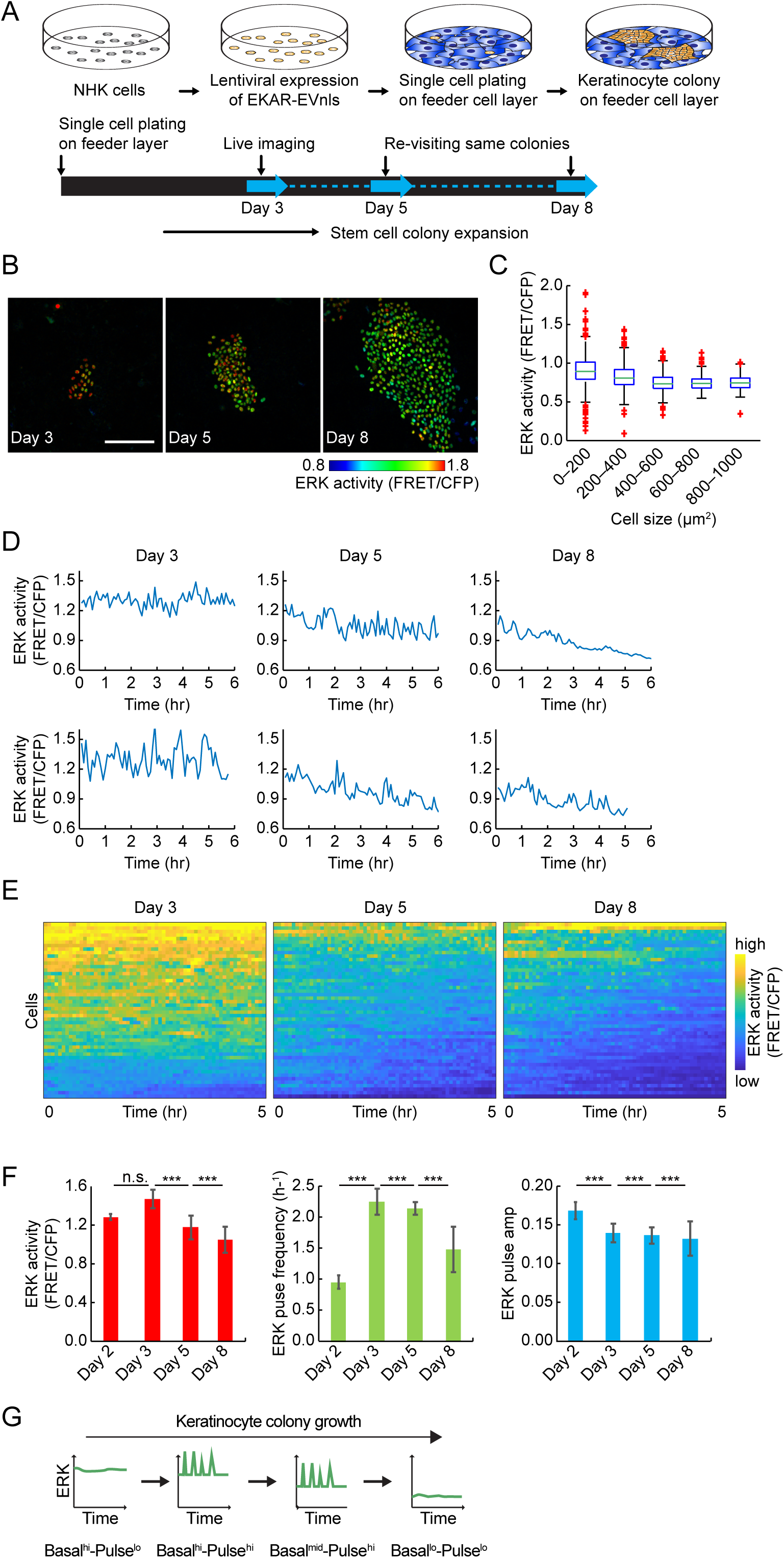
ERK activity dynamics transition in human epidermal stem cells. (*A*) Schematic of experimental settings. Primary human neonatal keratinocytes (NHKs) were lentivirally transfected with EKAR-EVnls ERK FRET sensor and allowed for colony formation on feeder cell layers. Same NHK colonies were repeatedly observed for live imaging. (*B*) Representative Images of NHKs expressing EKAR-EVnls on different days. The same colonies were observed at different days. Colours indicate ERK activity. Scale: 200 μm. (*C*) Box plots of single cell ERK activity grouped by cell size: mid-line, median; box, 25th to 75th percentiles; whiskers, lower and higher extremes; red crosses, outliers. (*n* = 3,581 cells). (*D*) Representative time-series of ERK activity on different days. (*E*) Heat-map of ERK activity over time for 50 cells ordered by descending mean ERK activity (FRET/CFP) over time. Colours indicate ERK activity. (*F*) Basal ERK activity (left), ERK pulse frequency (middle), and ERK pulse amplitude on different days indicated. Data are shown by mean ± s.e.m. (*n* = 542 cells for Day2, 3,323 cells for Day3, 11,527 cells for Day5, 37,320 cells for Day8 cells, two-tailed unpaired Student’s t-test; P values are indicated by *** P<0.001, n.s. = not significant (P>0.05). (*G*) Schematic representation of transitions in ERK activity dynamics comprising of basal ERK activity and its pulse activations during stem cell colony growth.

In order to examine ERK activity dynamics in detail we observed thousands of cells in growing keratinocyte colonies (Fig. 1 *D* and *E* and Movie S1; *n* = 3,323 cells for Day3, 11,527 cells for Day5, 37,320 cells for Day8). We found the ERK dynamics are characterised by pulse activations as well as its basal activity. We measured the quantitative features of ERK activity pulses as local peaks (Fig. S1*A*). We ruled out the possibility that the pulses were an imaging artefact by using a negative control FRET biosensor EKAREV-TA-nls, which has a mutation that results in loss of recognition by active ERK (Fig. S1*B*)(26). NHKs expressing EKAREV-TA-nls showed almost no pulse signals: only 5% of cells showed very rare (0.003 pulse/hr) pulses (Fig. S1*B*). The average duration of pulse activations in EKAR-EVnls expressing cells was 0.25 hr, which is consistent with that previously reported in immortalized epithelial cells(9, 10) (Fig. S1*C*). The average pulse-to-pulse interval was 1.52 hr. Notably, the histogram of interpulse intervals followed an exponential decay curve, which indicates that ERK pulses are stochastic rather than precisely timed events such as oscillations (Fig. S1*D*).

The frequency of pulses in NHKs ranged from zero to approximately 4.5 pulse/hr (Fig. 1*E* and Fig. S1*E*). The ERK pulse frequency was high (> 2.0 pulse/hr) on Day3 and Day5 of colony growth and subsequently reduced on Day8, while basal ERK activity gradually decreased (Fig. 1*F* and Fig. S1 *E* and *F*). In contrast, although ERK pulse amplitude decreased over the period the absolute reduction was very small (< 0.005 FRET/CFP; Fig. 1*F*), in line with previous studies in immortal cell lines(9, 10).

Detailed histogram analysis of basal ERK activity showed two peaks on Day3 (Fig. S1*F*, asterisks). The ERK activity in the lower peak matched that of the peak on Day5 (Fig. S1*F*). This suggested that the cell population with higher basal ERK activity on Day3 transited to the one with lower activity on Day5 rather than there being a gradual reduction in ERK activity among whole cell population.

We extended our analysis to include Day2 colonies (*n* = 542 cells). NHKs on Day2 showed significantly lower ERK pulse frequency and higher basal activity compared to older colonies (Fig. 1*F* and Fig. S1 *G-J*). We performed correlation analysis to determine whether basal ERK activity might influence ERK pulse activations. This revealed a moderate correlation on Day2, 3 and 8 (R = 0.23 – 0.65) but not on Day5 (R = 0.018) (Fig. S1*K*), strengthening the conclusion that ERK activity dynamics on Day5 are distinct from those on Day3.

These analyses show that individual human epidermal cells not only differ in basal ERK activity but also exhibit pulses of activation. Based on the analysis of individual cells in colonies at different times after plating, we propose that stem cells have a Basal^hi^-Pulse^lo^ ERK profile, then transit to Basal^hi^-Pulse^lo^, Basal^mid^-Pulse^hi^, and finally to Basal^lo^-Pulse^lo^ once they have undergone differentiation (Fig. 1*G*).

### Pulsatile ERK activations are associate with proliferation whereas pulse downregulation precedes differentiation

To confirm that transitions in ERK activity correlated with differentiation, we generated a fluorescent reporter of Involucrin, a marker gene that is upregulated in differentiating suprabasal epidermal cells. We used the previously characterized Involucrin promoter and intron sequence(27) to drive mCherry expression (Fig. 2*A*). Human epidermal stem cells lentivirally expressing the reporter were induced to differentiate by changing medium from low Ca^2+^ serum-free medium (KSFM)(28) to medium containing high Ca^2+^ (1.6 mM) or serum. NHKs expressing the Involucrin-mCherry reporter showed significant mCherry induction under both differentiation stimuli (Fig. 2 *B* and *C*).

**Fig. 2.**
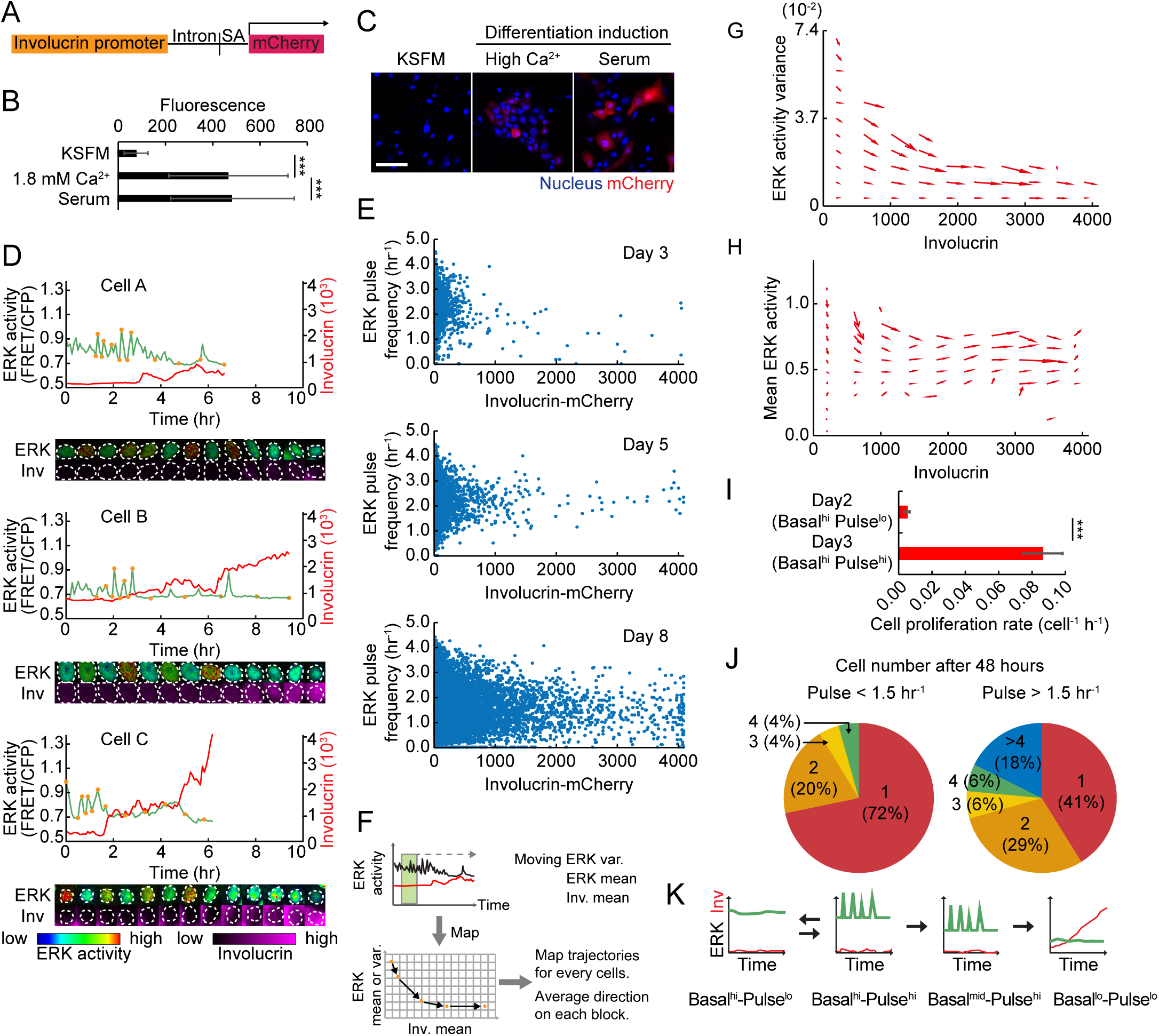
ERK pulse downregulation precedes commitment to differentiation. (*A*) Construction of Involucrin-mCherry reporter. SA: SV40 splice acceptor. (*B* and *C*) Involucrin-mCherry reporter expression in cells cultured under the indicated conditions. Data are mean ± s.d. (*n* = 50 cells each). Scale: 100 μm. (*D*) Representative time-series of ERK activity (black) and Involucrin-mCherry expression (red), and images of the cells at the time points indicated by the orange circles in each time-series. Images are shown by the indicated LUTs below. (*E*) Dot plot of ERK pulse frequency and Involucrin-mCherry expression in HNK cells on different days (*n* = 3,323 cells for Day3, 11,527 cells for Day5, 37,320 cells for Day8 cells). (*F*) Schematics of the methodology for constructing the phase diagram of ERK activity. ERK moving variance, and ERK and Involucrin moving mean levels are measured for each cell. Mapping of each ERK measure in relation to Involucrin generates a trajectory of the co-evolution of the two factors. (*G* and *H*) Phase diagram of ERK activity variance (*G*), and mean activity (*H*) against mean Involucrin expression obtained from *n* = 3,397 cells. Arrows indicate the direction of transition for each compartment. (*I*), The cell proliferation rate of NHK cells at ERK Basal^hi^-Pulse^lo^ state (Day2) and Basal^hi^-Pulse^hi^ state (Day3). Data are mean ± s.d. (*n* = 542 cells for Day2, 3,323 cells for Day3 cell). (*J*) Cell proliferation assay of single cells with ERK pulses lower (left) or higher (right) than 1.5 pulse/h. Cells were initially imaged at single cell state to measure their pulse levels, and then the same cells were observed after 48 hours. The number of cells at 48h and the proliferation fractions. (*K*) Schematic representation of modulations in ERK activation pulses during differentiation. Statistical significance was examined by two-tailed unpaired Student’s t-test; P values are indicated by *** P<0.001.

ERK activity and differentiation were simultaneously monitored by co-expression of the EKAR-EVnls and Involucrin reporters in individual keratinocytes. We found that ERK pulses were downregulated coincident with the onset of Involucrin expression (Fig. 2*D*) while cells that maintained low or high Involucrin expression showed stable ERK activity profiles (Fig. S2 *A* and *B*). As expected, during NHK colony growth, the number of cells expressing Involucrin-mCherry increased and they tended to have lower ERK basal activity and pulse frequencies (Fig. 2*E* and Fig. S2*C*). The reduction in pulse frequency, however, did not show a strong correlation with Involucrin-mCherry expression levels in individual cells. (Fig. 2*E*).

To further dissect the change in ERK activity associated with differentiation we analysed the trajectories of ERK activity (mean and variance) and Involucrin expression over time (Fig. 2*F*). We observed co-evolution of ERK activity variance and Involucrin expression, showing that there is a strong tendency for undifferentiated cells to downregulate pulse frequencies coupled with differentiation (Fig. 2*G*). In contrast there was a gradual convergence of basal ERK activity towards a low level as differentiation proceeded (Fig. 2*H*). This difference suggests that ERK pulses and mean levels are subject to distinct regulatory mechanisms, and that the downregulation in ERK pulses has a role in switching stem cells to differentiated cells. The trajectory of ERK dynamics in Basal^hi^-Pulse^hi^ (Day3 after plating) cells showed that the cells tend to decrease pulse frequency and increase mean activity, which will lead to Basal^hi^-Pulse^lo^ ERK activity (Fig. S2*D*). This suggests that the ERK dynamics is partially reversible in the early stage while it is irreversible once they are committed to differentiation.

Epidermal keratinocytes not only transition from the stem cell compartment to the differentiation compartment but can also transition between cell division and quiescence(14). To determine whether ERK pulse activations were associated with cell division, we followed the fate of Basal^hi^-Pulse^lo^ cells (Day2 after plating) and Basal^hi^-Pulse^hi^ (Day3 after plating) that did not subsequently express Involucrin (Fig. 1*E*). Basal^hi^-Pulse^hi^ cells had a high probability of dividing, whereas Basal^hi^-Pulse^lo^ cells did not (Fig. 2*I*).

We also recorded the ERK pulse frequency of individual cells and whether or not they subsequently divided (within 48 hours). We found that 70% of cells with low ERK pulse frequency (< 1.5 pulse/h) remained as single cells (Fig. 2*J*, left), whereas 60% of cells with high ERK pulse frequency (> 1.5 pulse/h) proliferated, giving rise to 2 or more cells during the recording period (Fig. 2*J*, right). This indicates that cells with a pulsatile ERK profile are more likely to divide than cells with a stable-high ERK profile.

These results, combined with our observations of cells expressing Involucrin-mCherry suggest that stem cells transition from Basal^hi^-Pulse^lo^ to Basal^hi^-Pulse^hi^ when proliferating, and that a subsequent transition to Basal^mid^-Pulse^hi^ is associated with differentiation, leading to Basal^lo^-Pulse^lo^ ERK activity in differentiated cells (Fig. 2*K*).

### ERK pulse modulation by terminal differentiation stimuli

We next tested the effect of different differentiation stimuli on ERK activity dynamics. Integrin-mediated adhesion maintains the stem cell state via ERK signalling(29). We found by siRNA-mediated knockdown that reduced β1-integrin expression strongly induced ERK pulses in the whole cell population (Fig. 3 *A* and *B* and Fig. S4 *A*-*C*).

**Fig. 3.**
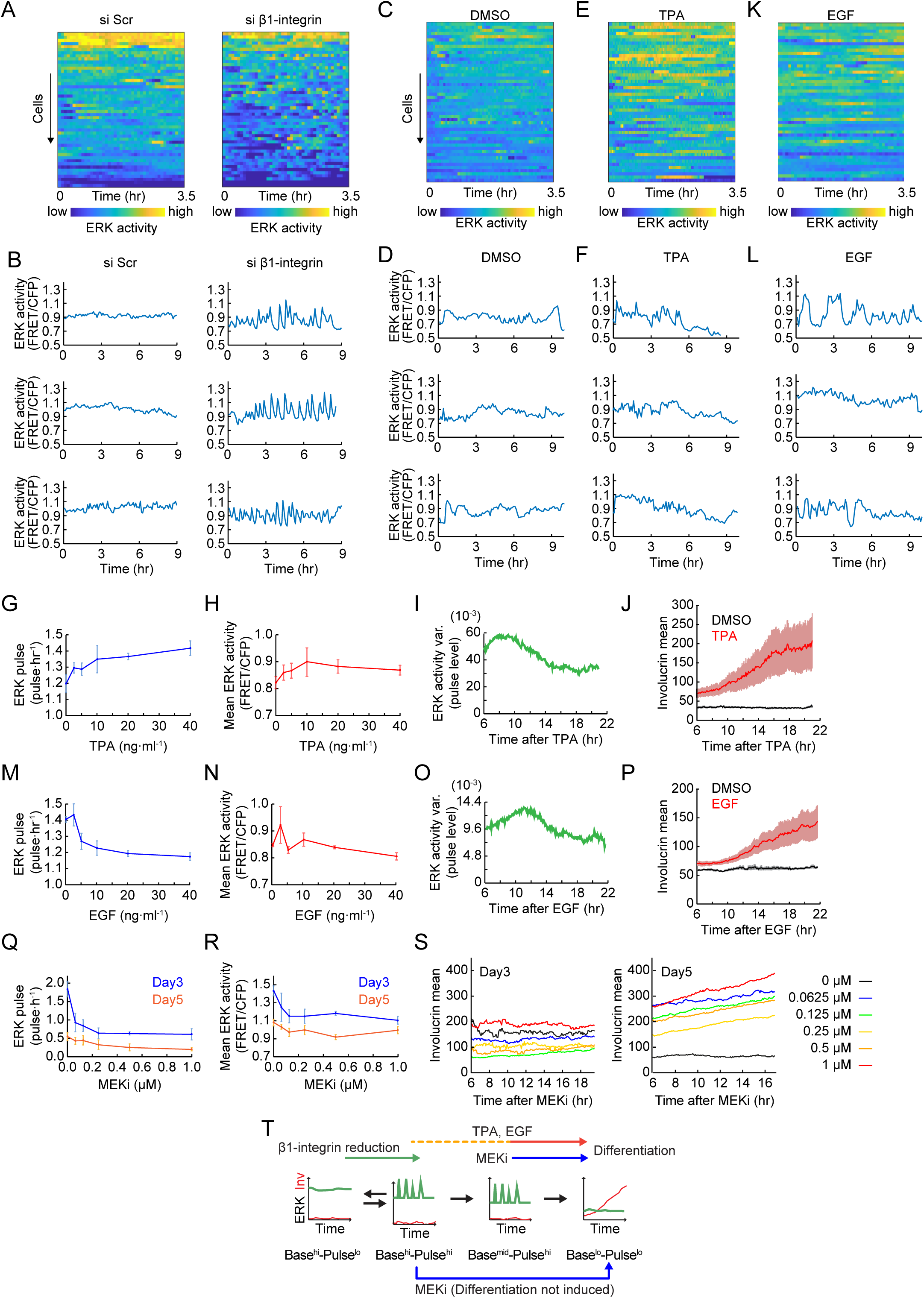
ERK pulse dynamics in induced cell differentiation. (*A* and *C* and *E* and *K*) Heat-maps of ERK activity over time for 50 cells ordered by descending overall mean ERK activity in each culture condition. Colours indicate ERK activity. (*B* and *D* and *F* and *L*) Representative time-series of ERK activity in cells treated with the indicated reagents. (*G* and *H*) Dose dependency of ERK pulse frequency (*G*) and mean activity (*H*) on TPA treatment. Data are shown by mean ± s.d. (*n* = 1,071 ± 27 cells for each condition). (*I* and *O*) ERK activity variance over time (see *Materials and Methods*) in cells treated with 10 ng/ml TPA (*I*) or 10 ng/ml EGF (*O*). (*J* and *P*) Involucrin expression over time in cells treated with DMSO, 10 ng/ml TPA (*J*), or 10 ng/ml EGF (*P*). Data are shown as mean ± s.d. (*M* and *N*) Dose dependency of ERK pulse frequency (*M*) and mean activity (*N*) on EGF treatment. Data are shown by mean ± s.d. (*n* = 937 ± 158 cells for each condition). (*Q* and *R*) Dose dependency of ERK pulse frequency (*Q*) and mean activity (*R*) on MEK inhibitor PD0325901 treatment in NHKs cultured for 3 or 5 days. Data are shown by mean ± s.d. (*n* = 670 ± 149 cells for each condition on Day3, and 885 ± 438 cells for each condition on Day5). (*S*) Involucrin expression over time in cells treated with the indicated dose of MEK inhibitor, PD0325901 on Day3 or Day5 NHKs. Data are shown as mean ± s.d. (*n* = 670 ± 149 cells for each condition on Day3, and 885 ± 438 cells for each condition on Day5). (*T*) Schematic representation of the effects of differentiation stimuli an ERK activity dynamics.

In contrast, blocking cell-cell adherens junctions and desmosomes by Ca^2+^ depletion of FAD medium had little effect on ERK pulses (Fig. S4 *D*-*H*, 1.15 pulses/hr vs 1.06 pulses/hr). Consistent with this, Involucrin expression was not affected by the inhibition of intercellular adhesion (Fig. S4*I*)(30). These results indicate that modulation of cell-substrate interaction plays a more significant role than Ca^2+^-mediated cell-cell interaction, and provides further experimental evidence of the transition in temporal ERK patterns during differentiation (Fig. 1*F*).

12-O-Tetradecanoylphorbol-13-acetate (TPA) is known to stimulate Involucrin expression(31) and also increases ERK activity(32). When cells were stimulated with 10 ng/ml TPA, they exhibited highly pulsatile ERK activity (Fig. 3 *C*–*F*). Overall ERK pulse frequency increased in a dose dependent manner up to 40 ng/ml TPA (Fig. 3*G*), while mean ERK levels peaked at 10 ng/ml (Fig. 3*H*). The time-course analysis revealed that TPA induction of ERK pulses was transient, peaking at 9 hr after the start of treatment (Fig 3*I*). As before (Fig. 2 *G* and *K*), the onset of Involucrin expression coincided with the downregulation of ERK pulses (Fig. 3*J*).

Like TPA, epidermal growth factor (EGF) transiently enhanced ERK pulse levels (Fig. 3 *K* and *L*), although, in contrast to TPA, EGF decreased overall ERK pulse frequency in a dose dependent manner without affecting ERK mean levels (Fig. 3 *L* and *M*). ERK pulses peaked at 11 hr after EGF treatment and then showed a significant decrease, again coincident with the onset of Involucrin expression (Fig. 3 *O* and *P*). Thus TPA and EGF had different effects on overall ERK pulse frequency (Fig. 3 *G* and *M*) but the time course of changes is similar and in both cases ERK was downregulated when cells expressed Involucrin (Fig. 3 *I* and *J* and *O* and *P*). This leads us to speculate that the downregulation of ERK pulses triggers differentiation (Fig. 2*K*).

When human keratinocytes were treated with a MEK inhibitor PD0325901 (MEKi), they exhibited a dose-dependent reduction in both ERK pulse frequency and basal levels (Fig. 3 *Q* and *R* and Fig. S4 *A*-*D*). We tested the inhibitor on different days after plating cells: Day3 (correlating with Basal^hi^-Pulse^hi^; Fig. 1*F* and Fig. S1 *E* and *F*) and Day5 (correlating with Basal^mid^-Pulse^hi^, Fig. 1*F* and Fig. S1 *E* and *F*) (Fig. S4 *E* and *F*). Although on both days MEKi induced the Basal^lo^-Pulse^lo^ state, differentiation was only induced on Day5 (Fig. 3*S*). This suggests that the order of transitions in ERK dynamics shown in Fig. 2*K* must be followed for differentiation to occur.

We conclude that three distinct differentiation stimuli – reduced integrin-mediated adhesion, TPA and EGF – all trigger ERK pulses and subsequent ERK downregulation, whereas inhibition of MEK reduces ERK basal levels directly. Furthermore, cells initiate differentiation by transiting through the Basal^mid^-Pulse^hi^ state (Figs. 1*F* and 2*K* 3*T*).

### Regulation of ERK mean and pulsatile activity by protein phosphatases

One likely mechanism by which ERK basal activity and pulses are controlled is via negative feedback regulation by protein phosphatases(33). We therefore examined the effects of DUSP6 and DUSP10, key members of the protein phosphatase network that acts as a commitment switch in human epidermal stem cells(23). By live imaging cells in which overexpression of each DUSP was induced by doxycycline, we found that DUSP6 reduced ERK pulses without changing basal ERK activity (Fig. 4*A*), while DUSP10 downregulated basal ERK levels without changing ERK pulses (Fig. 4*B*). This indicates that mean ERK levels and ERK pulses are independently regulated by different phosphatases.

**Fig. 4.**
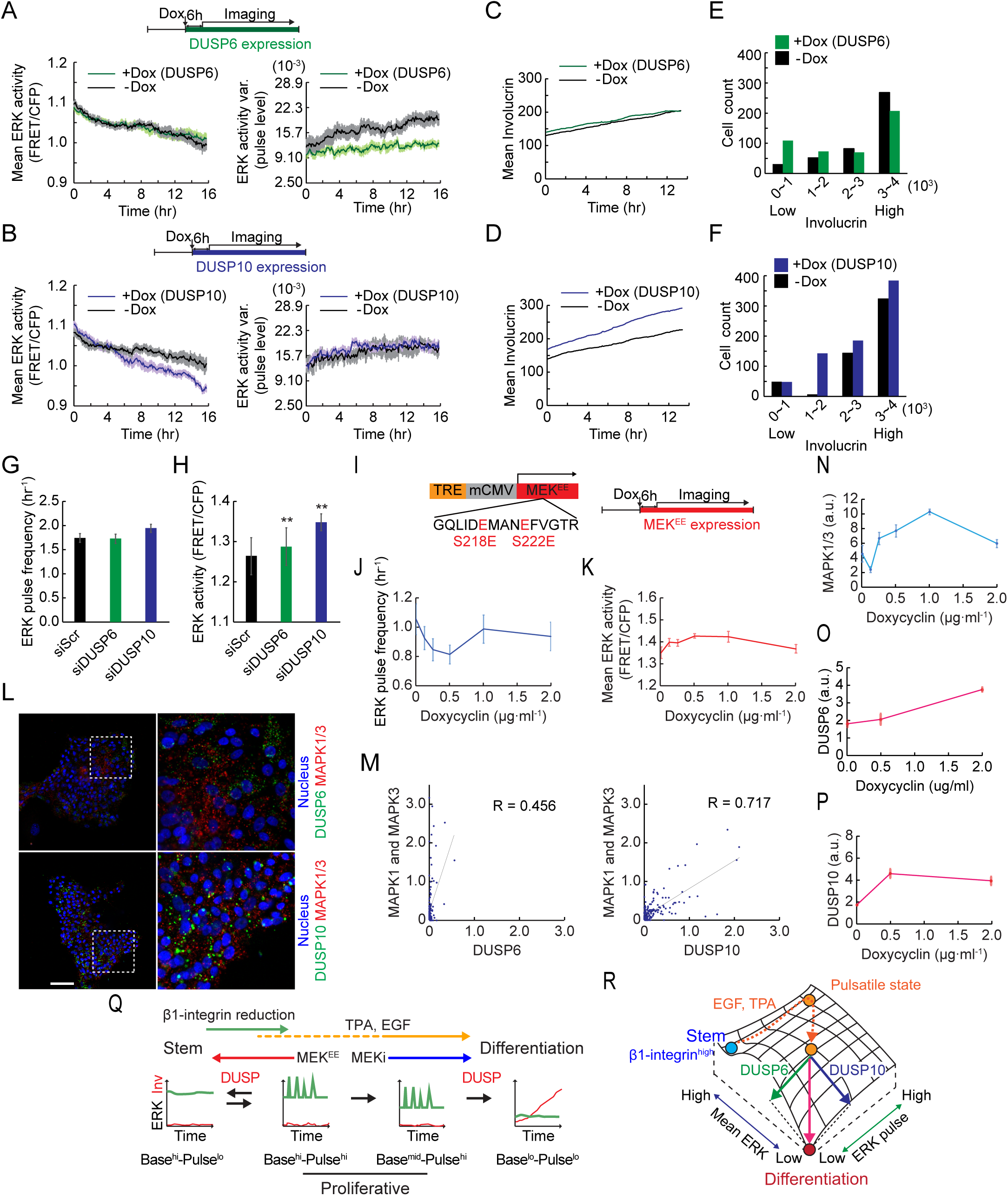
ERK pulse and mean levels are independently regulated by DUSP6 and DUSP10. (*A* and *B*) Mean (left) and fluctuation levels (right, see *Materials and Methods*) of ERK activity in NHKs treated with 1 µg/ml doxycyclin (green or purple) or vehicle (black). DUSP 6 (*A*), or DUSP10 (*C*), was induced by 1 µg/ml doxycycline treatment. Data are shown by mean ± s.e.m. (*n* = 1,220 doxycyclin-treated cells and 1,261 vehicle-treated cells for (*A*), *n* = 1,224 doxycyclin-treated cells and 1,005 vehicle-treated cells for (*B*)). (*C* and *D*) Involucrin expression over time in NHKs treated with 1 µg/ml doxycyclin (green or purple) or vehicle (black). DUSP 6 (*C*), or DUSP10 (*D*), was induced by the doxycycline treatment. (*E* and *F*) Cell number increase in the four bins of Involucrin-mCherry expression level 18.5 hour after doxycycline-induced DUSP6 and DUSP10 expression (green and purple, respectively) or vehicle (black) treatment. (*G* and *H*) ERK pulse frequency (*G*) and mean ERK activity (*H*) in NHKs treated with scramble siRNA, siRNA against DUSP6, or siRNA against DUSP10. Data are shown by mean ± s.e.m. (*n* = 5,701 siScr cells, 4,335 siDUSP6 cells, and 4,346 siDUSP10 cells, two-tailed unpaired Student’s t-test; P values are indicated by ** P<0.01). (*I*) Construction of plasmid for doxycyclin-dependent expression of constitutive active MEK1 (MEK^EE^). For MEK^EE^ induction, cells were treated with 0.125 - 2.0 µg/ml doxycyclin. (*J* and *K*) ERK pulse frequency (*J*) and mean ERK activity (*K*) of NHKs treated with indicated dose of doxycycline to induce MEK^EE^ expression. Data are shown by mean ± s.e.m. (*n* = 1,140 ± 494 cells for each condition). (*L*) Representative images of RNA In Situ Hybridization of DUSP6 (above, green), DUSP10 (below, green), and MAPK1 and MAPK3 (red). The right images show close views of the white-dotted squares in the left images. Scale: 100 μm. (*M*) Dot plots and correlation analysis of DUSP6 (left) or DUSP10 (right), and MAPK1 and MAPK3 In Situ Hybridization signals in NHKs cultured for 5 days. Lines indicate regression lines and values are Spearman’s rank correlation coefficient. Data are shown by mean ± s.e.m (*n* = 66 cells for DUSP6 and 170 for DUSP10). (*N*-*P*) Quantification of RNA In Situ Hybridization signals of MAPK1 and MAPK3 (*N*), DUSP6 (*O*), and DUSP10 (*P*) in NHKs treated with indicated doses of doxycycline for MEK^EE^ expression. Data are shown by mean ± s.e.m. (*n* = 202 ± 82 cells for (*N*), 219 ± 115 cells for (*O*), 174 ± 44 cells for (*P*)) (*Q*) Schematic representation of the molecular regulations of ERK dynamics transition. (*R*) Schematic representation of the regulation of ERK mean and pulse level by DUSP6 and DUSP10 during differentiation.

Whereas DUSP10 overexpression strongly stimulated Involucrin expression during the recording period, DUSP6 did not have a significant effect (Fig. 4 *C* and *D*). However, by binning cells into groups of low/intermediate/high Involucrin levels, we found that DUSP6 and DUSP10 induction had different effects on each population. DUSP6 increased the proportion of cells with low Involucrin expression (Fig. 4*E*). In contrast, DUSP10 increased the proportion of cells with high Involucrin expression (Fig 4*F*). This suggests the intriguing possibility that DUSP6-mediated ERK pulse downregulation promotes the initiation of differentiation whereas DUSP10-mediated downregulation of mean ERK activity promotes and stabilizes post-commitment differentiation. This is consistent with the finding that DUSP6 is transiently upregulated on commitment, while DUSP10 upregulation is more sustained(23).

We also tested siRNA-mediated knockdown of DUSP6 and DUSP10(23). Reduced DUSP6 expression did not have a significant effect on ERK pulse frequency or basal activity (Fig. 4 *G* and *H*). However, reduction in DUSP10 expression led to increased basal ERK activity while maintaining pulse frequency (Fig. 4 *G* and *H*). This confirms that DUSP10 controls basal ERK activity.

As a further means of perturbing ERK activity we lentivirally transfected HNK cells with a constitutively active form of MEK1 (MEK^EE^). MEK1 lies immediately upstream of ERK in the ERK signalling cascade and can over-ride the differentiation stimulus of reduced β1-integrin signalling in keratinocytes (34). Moderate induction of MEK^EE^ expression with 0.5 µg/ml doxycycline (Fig. 4*I*) significantly decreased ERK pulse frequency (Fig. 4*J*) and increased basal ERK activity (Fig. 4*K*), promoting the transition from Basal^hi^-Pulse^hi^ to Basal^hi^-Pulse^lo^ ERK.

### Transcriptional control of ERK, DUSP6 and DUSP10

Previous studies have demonstrated interactions between DUSP6, DUSP10 and ERK at the transcriptional level in addition to the post-translation level(23, 35-37). We therefore compared transcripts of MAPK3 and MAPK1, which code ERK1 and ERK2 proteins, together with DUSP6 and DUSP10, in individual cells using RNA fluorescence *in situ* hybridization (Fig. 4*L*). MAPK1/3 expression was significantly correlated with DUSP10 and DUSP6 expression (Fig. 4*M*). In addition, the level of MEK^EE^ induction that decreased ERK pulse frequency and increased basal ERK activity (Fig. 4 *J* and *K*) also increased expression of MAPK1, MAPK3, DUSP6 and DUSP10 transcripts (Fig. 4 *N*-*P*). We conclude that ERK activity in keratinocytes is subjected to both transcriptional and post-transcriptional regulation and that there are compensatory mechanisms to prevent excessive upstream stimulation of ERK activity (Fig. 4*R*).

Together, β1-integrin, EGF and its downstream effectors underlie the ERK dynamics transitions to achieve different cellular outcomes (Fig. 4*Q*). DUSP expressions lead to Pulse^lo^ states whether the basal ERK activity is high or low. The Base^hi^-Pulse^lo^ state could be the result of transcriptional upregulation of MAPK1/3 that potentially cancel the effect of DUSP10 to reduce basal ERK activity. During differentiation, DUSP6 and DUSP10 independently downregulate ERK pulses and basal activity (Fig. 4*R*). The loss of correlation between ERK basal activity and pulse frequency in the Base^mid^-Pulse^hi^ state (Fig. S1*K*) might be important for the independent downregulations of the two ERK features.

### Patterning of keratinocytes with different ERK kinetics in response to substrate topography

We have previously reported that human epidermal stem cells have a patterned distribution in skin(38), leading us to predict that ERK dynamics would be spatially regulated. To examine this, we plated NHKs co-expressing a cytoplasmic ERK sensor (EKAR-EVnes) and Involucrin-mCherry on collagen-coated undulating polydimethylsiloxane (PDMS) substrates that mimic the topography of the human epidermal-dermal junction(23, 39). As in human epidermis, stem cells that express high β1-integrin and DUSP6 levels cluster at the tips of the features, whereas DUSP10 expression is more uniformly distributed(23). Once the cells had formed a confluent multi-layered sheet they were subjected to live cell imaging (Fig. 5*A*). As reported previously(39), Involucrin-positive keratinocytes accumulated at the base of the features (troughs), while Involucrin-negative cells accumulated on the tips (Fig. 5 *B* and *C*).

**Fig. 5.**
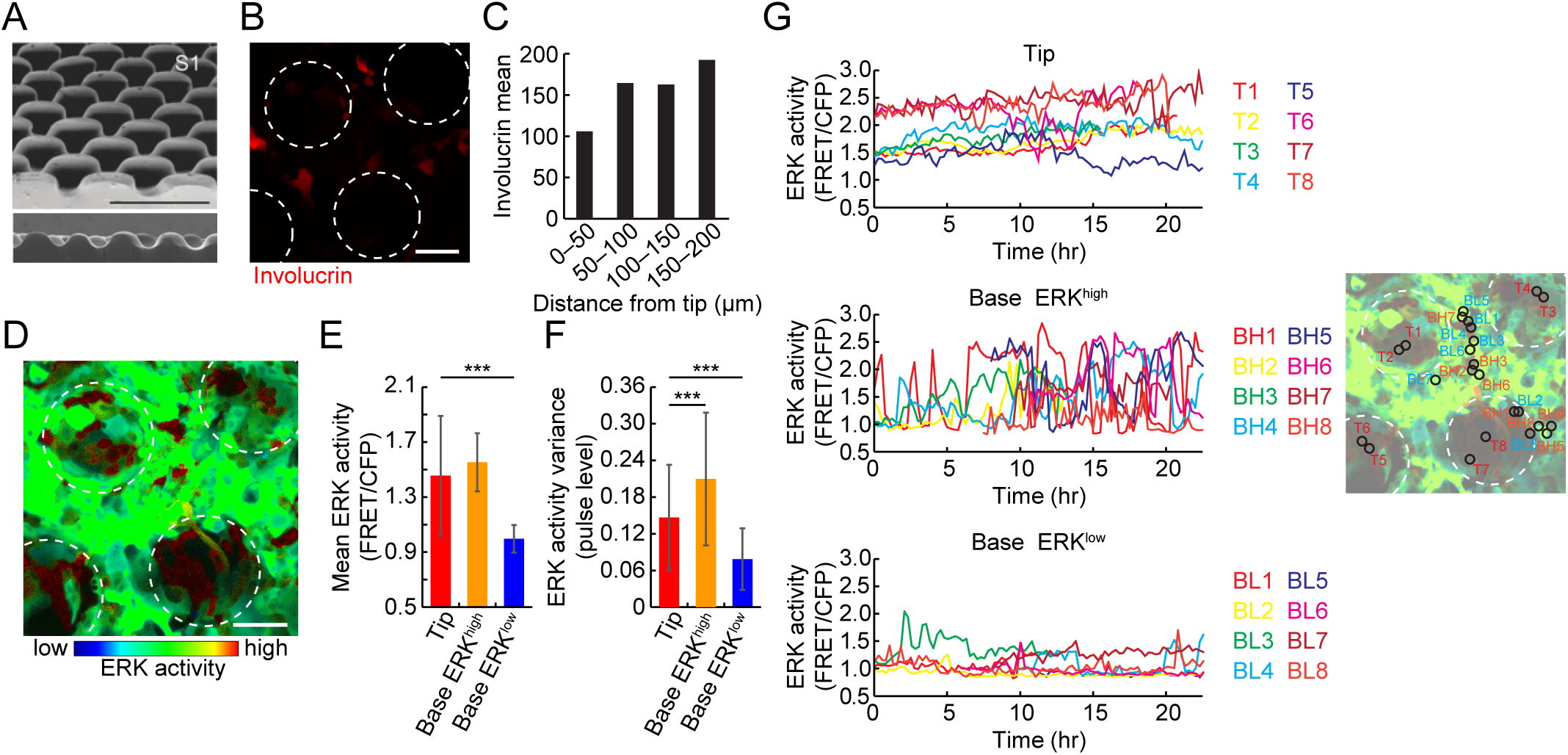
Spatial segregation of different ERK time-series patterns on topographical substrates. (*A*) Overview (above) and cross section (below) SEM images of the patterned PDMS substrate. Reproduced from a previous publication(39). Scale: 400 μm. (*B*) Involucrin-mCherry expression on the patterned PDMS substrate. White dotted circles: tips of the substrate. Scale: 100 μm. (*C*) Mean Involucrin-mCherry expression in regions at different distances from the tips. (*D*) Representative image of ERK activity in human keratinocytes cultured on the PDMS substrate. Colours indicate ERK activity. White circles: tips of the substrate. Scale: 100 μm. (*E* and *F*) ERK activity mean (*E*), and pulse level (*F*), for cells cultured on patterned substrate. Data are shown as mean ± s.d. (*n* = 50 Tip cells, 27 Base ERK^high^ cells, and 23 Base ERK^low^ cells, two-tailed unpaired Student’s t-test; P values are indicated by *** P<0.001). (*G*) Representative time-series of human keratinocytes on tip regions (above) and those in base regions (troughs) with high (middle) and low (below) ERK activity. Eight time-series are shown for each group. The locations of the cells are mapped on the image to the right.

We observed a patterned distribution of ERK activity on the substrates. Cells on the tips had higher basal ERK activity and lower ERK pulse frequencies than cells in the troughs (Fig. 5 *D* and *E* and Movie S2). Tip-located cells were also less motile (Fig. 5 *D*-*G* and Movie 2), consistent with the high β1-integrin expression and low motility of epidermal stem cells(39). Conversely, cells in the troughs and sides of the substrates had with low-stable ERK activity or pulsatile activity (Fig. 5 *D*-*G* and Movie 2). Those cells in the troughs with low Involucrin expression had a higher level of ERK pulsatile activity than other cells (Fig. 5 *E*-*G*).

We conclude that on a 3D topography that mimics the human epidermal-dermal interface, cells with distinct patterns of ERK activity were differentially localized. The tip “stem cell niche” regions were occupied by cells with a stable-high ERK activity(40), while the base regions were occupied by cells with pulsatile ERK patterns or cells with low-stable activity that underwent differentiation.

### ERK pulse kinetics are preserved in mouse epidermis

By live imaging of human epidermal cells we found that downregulation of pulsatile ERK activity preceded terminal differentiation and that ERK activity was patterned according to the location of stem cells and differentiated cells. To test whether this was also the case in living tissue, we generated mice that express both EKAR-EVnls and an Involucrin-tdTomato reporter(41). In interfollicular epidermis of mouse, as in human skin, the stem cells reside in the basal layer and differentiating cells occupy the suprabasal layers (Fig. 6 *A* and *B*). We imaged two different regions of the skin, in which epidermal cells have distinct patterns of proliferation and differentiation: the ear(42) and tail(43).

**Fig. 6.**
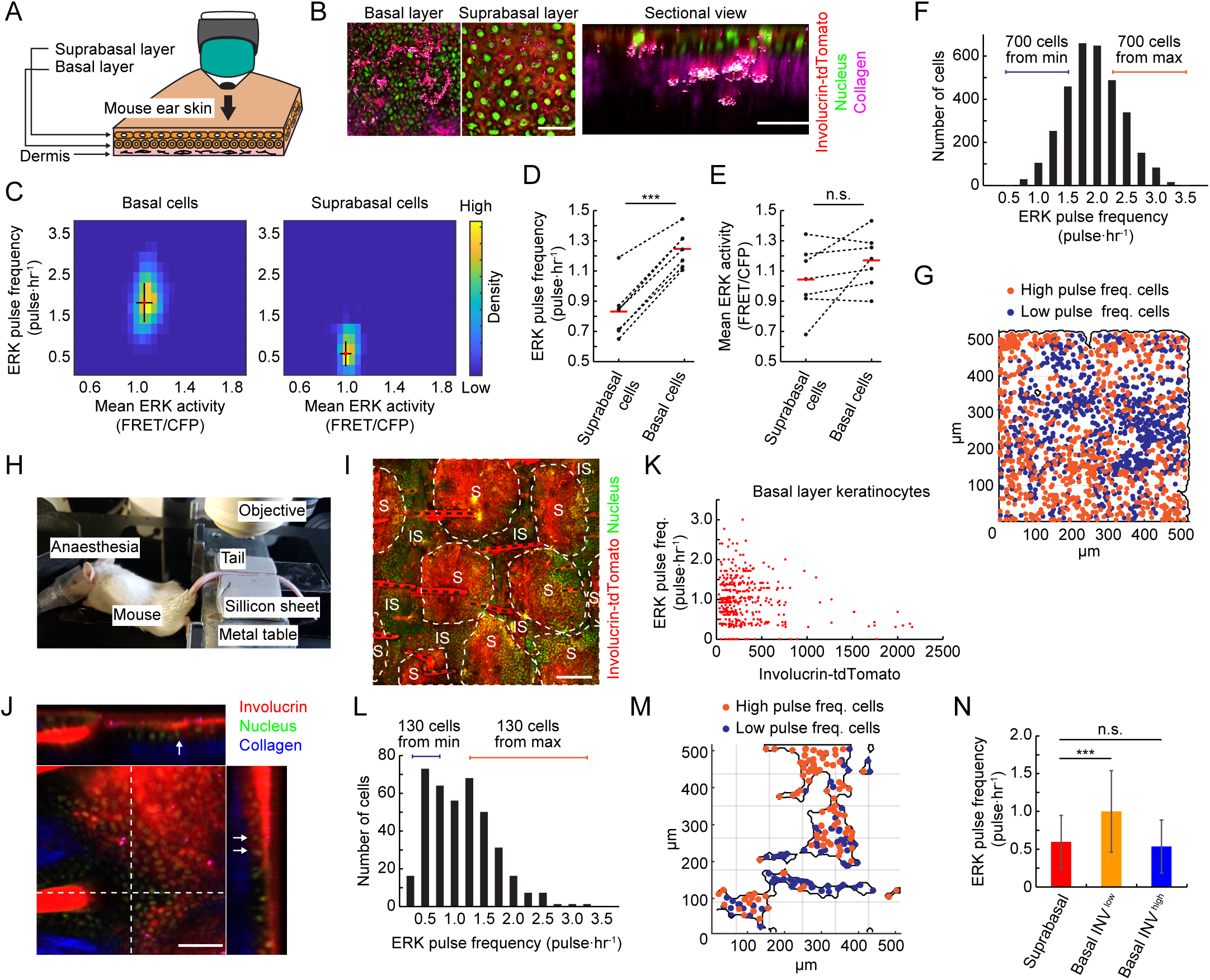
ERK pulse kinetics is preserved in live mouse epidermis. (*A*) Schematic of in vivo observation of mouse epidermis. Ear skin of mouse epidermis expressing EKAR-EVnls and Involucrin-tdTomato was observed by multiphoton microscopy. (*B*) Representative images of epidermis in mouse expressing EKAR-EVnls and Involucrin-tdTomato. Collagen (magenta) was visualized by second harmonic generation microscopy. Sectional view image was reconstructed by z-stack images. Scale: 100 μm. (*C*) 2D histogram of ERK pulse frequency and mean activity in basal (left) and suprabasal (right) layer cells. Mean and s.d. are shown by red dot and black lines, respectively (*n* = 3,238 basal cells and 352 suprabasal cells). (*D* and *E*) ERK pulse frequency (*D*), and mean ERK activity (*E*), in the basal and suprabasal layer cells of 7 mice. Mean is shown by red bars. (*F*) Distribution of ERK pulse frequencies of cells in the basal layer of mouse ear skin. (*n* = 3,238 cells). (*G*) Mapping of basal layer keratinocytes showing the 700 least pulsatile (blue) and 700 most pulsatile (orange) cells, as indicated in (*F*). (*H*) In vivo imaging of mouse tail epidermis. The anaesthetized mouse was placed on the heater and the tail skin was stabilized between soft silicon gum sheets. (*I*) Projection XY view of tail epidermis of mouse expressing EKAR-EVnls and Involucrin-tdTomato. S: scale epidermis, IS: interscale epidermis, black dotted lines: hair. Scale: 200 μm. (*J*) Orthogonal views of tail epidermis of the mouse expressing EKAR-EVnls and Involucrin-tdTomato. Arrows indicate basal layer cells expressing Involucrin. (*K*) Dot plot of ERK pulse frequency and Involucrin-tdTomato expression in individual cells (*n* = 391 cells). (*L*) Distribution of ERK pulse frequencies of cells in the basal layer of mouse tail skin (*n* = 391 cells). (*M*) Mapping of basal layer keratinocytes showing the 130 least pulsatile (blue) and 130 most pulsatile (orange) cells as indicated in (*L*). (*N*) Mean ERK pulse frequencies in the indicated tail epidermal cells. Data are shown by mean ± s.d. (*n* = 318, 374, 19 cells). Statistical significance was examined by two-tailed unpaired Student’s t-test; P values are indicated by *** P<0.001, n.s. = not significant (P>0.05).

In the skin of anaesthetized mice the boundary between the epidermis and the underlying dermis could readily be visualized by second harmonic generation (SHG) microscopy of collagen. Differentiating cells expressed tdTomato, and all cell nuclei expressed EKAR-EVnls (Fig 6 *B* and *I* and *J*). Time lapse observation of ERK activity revealed that basal keratinocytes had significantly higher ERK pulse levels than differentiating, tdTomato-positive suprabasal keratinocytes (Fig. 6*C* and S5 *A* and *B*).

Quantitative analysis of ear epidermis revealed that there was a greater difference in ERK pulse levels (1.88 pulse/hr vs 0.64 pulse/hr) than ERK mean levels (1.12 pulse/hr vs 1.05 pulse/hr) between basal and suprabasal cells (Fig. 6*C*). Observation of multiple mice showed consistent differences in ERK pulse levels between the basal and suprabasal layers, whereas the differences in ERK mean levels were relatively limited and highly variable among the seven mice examined (Fig. 6 *D* and *E*). This suggests a more significant role of ERK pulses than basal activity in the epidermis of living mice.

Ear epidermis is organized into columns of differentiated cells arranged above groups of basal cells, which have been referred to as Epidermal Proliferative Units (EPUs)(42, 44). The width of an ear EPU is approximately 25 μm diameter, with approximately 8 – 10 cells in the basal layer. We noticed a high variance in ERK pulse frequencies in the basal layer of ear epidermis and therefore mapped the distribution of basal cells with high or low ERK pulse levels (Fig. 6*F* and Fig. S5 *G* and *H*). This revealed that cells with low ERK pulse levels were clustered (Fig. 6*G* and Fig. S5*C*). The cluster sizes were estimated to be about 50 μm (Fig. S5*C*), which is similar to the reported EPU size(42, 44). This segregation of cell clusters with different ERK dynamics is reminiscent of spatial segregation of cells with different ERK profiles on substrates mimicking human epidermis (Fig. 5*D*).

In the interfollicular epidermis of mouse tail skin there are two distinct programmes of terminal differentiation: scale (parakeratosis) and interscale (orthokeratosis)(43). The scale forms postnatally and it has been speculated that postnatal expansion is limited by a subset of keratinocytes that express Involucrin in the basal layer of the interscale(45). This led us to predict that the spatial distribution of basal cells with pulsatile ERK activity would differ between tail and ear skin, and also enabled us to monitor Involucrin-positive cells in the basal layer of tail epidermis. We imaged the tails of mice expressing both EKAR-EVnls and the Involucrin-tdTomato reporter (Fig. 6 *H* and *I* and Movie S3) and confirmed that some basal layer keratinocytes expressed Involucrin (Fig. 6*J*, arrows). In vivo time-lapse imaging of the mice revealed that ERK pulses were significantly reduced in basal layer keratinocytes expressing Involucrin compared to Involucrin-negative basal cells (Fig. 6*K*).

In the tail epidermal basal layer, ERK Pulse^hi^-Involucrin^lo^ and ERK Pulse^lo^-Involucrin^hi^ cells were mostly intermingled (Fig. 6 *L* and *M*). In contrast to ear epidermis, there was no significant clustering of low or high ERK pulse cells (Fig. 6*M* and Fig. S5*D*). We found that the average ERK pulse frequency in Involucrin^hi^ basal cells was comparable to that in suprabasal differentiated cells (Fig. 6*N*). This rules out the possibility that the reduced ERK pulses in Involucrin expressing cells of the ear are an artefact of imaging different layers of skin. It also confirms the strong coupling of reduced ERK pulses and Involucrin expression.

Together, our results indicate that ERK pulses are robustly coupled with cell fate, both in cultured human epidermal stem cells and in vivo mouse epidermal cells. The clustered localization and Basal^hi^-Pulse^lo^ stem cells in culture (Fig. 5*D*) and in mouse skin (Fig. 6*G* and Fig. S5*C*) is fully compatible with the idea that stem cells reside in specific niches that modulate key signalling pathways(46).

## Discussion

We have demonstrated that basal and pulsatile ERK activation dynamically regulate epidermal stem cell fate. Previous studies have shown that ERK plays a key role in exit of embryonic stem cells from pluripotency and in lineage specification(47, 48) but the significance of cell-to-cell variation in ERK activity has been unclear. We show that ERK pulse frequencies are regulated independently of basal ERK activity, and that dynamic ERK activity is a feature of both cultured human epidermis and the epidermis of living mice.

ERK pulses have been reported in multiple cell types in relation to cell proliferation and tissue morphogenesis(9, 10, 13, 49, 50). Our research demonstrates that ERK pulses play significant roles in stem cell fate regulation and raises the intriguing possibility that pulse-mediated changes in cell fate are conserved in multiple tissues and organisms. Indeed, in vivo imaging approaches have shown pulsatile ERK activity in mouse mammary gland and intestine(10, 51, 52).

Our finding of independent regulation of ERK pulse and mean levels by DUSP6 and DUSP10 indicates that DUSP6 and DUSP10 operate as independent negative feedback loops to achieve different ERK activity profiles and different cellular outcomes. We also observed patterning of cells with different types of ERK activity on culture substrates that mimic the topography of the epidermal-dermal junction, consistent with the different patterns of DUSP6 and DUSP10 expression(39). ERK activity was subject to both transcriptional and post-transcriptional regulation and subject to compensatory mechanisms to prevent excessive stimulation of ERK on constitutive activation of MEK1. In mouse epidermis there was a clear difference in ERK pulse frequencies, but not basal levels, in the stem and differentiated cell layers. This suggests that ERK pulse modulation is a strategy for cells to switch states in vivo as well as in vitro. Differences in the spatial segregation of cells with different ERK profiles between ear and tail skin are likely to reflect differences in the architecture of the stem cell niche.

Fluctuations of signalling pathways are increasingly recognized as key determinants for tissue development(6). Multifaceted features of those fluctuations, such as phase, frequency, and amplitude, provide potentially different outputs in terms of cell fate. It will now be of significant interest to explore whether other signalling pathways show pulsatile activity in the epidermis and, if so, how they interact with ERK.

## Materials and Methods

### Reagents

Hoechst 33342 was obtained from Molecular Probes, TPA (phorbol-12-myristate 13-acetone), EGF and PD0325901 were obtained from Sigma. RNAscope probes against human MAPK1, MAPK3, DUSP6 and DUSP10 were purchased from Advanced Cells Diagnostics (Probe-Hs-MAPK1, catalogue number: 470741-C2, Probe-Hs-MAPK3, catalogue number: 470731-C2, Probe-Hs-DUSP6, catalogue number: 405361, Probe-Hs-DUSP10, catalogue number: 583311).

### Constructs

pCSII-EKAREV-nls and pCSII-EKAREV-nes were previously described (53). EKAREV-TA-nls, negative control ERK FRET biosensor, is a kind gift from Dr. Eishu Hirata at Cancer Research Institute of Kanazawa, Japan, and was subcloned into pCSII vector. To construct pLenti-Involucrin-mCherry, the sequence comprising the full length 3.7kb Involucrin promoter, Involucrin intron, and SV40 splice donor and acceptor (S_D_S_A_) was subcloned from a previously-reported beta-galactosidase reporter(54) into pLenti backbone, and mCherry was tagged to the carboxy (C) terminus of the promoter sequence. Doxycycline-inducible DUSP expression plasmids (pCW57-DUSP6 and pCW57-DUSP10) were previously described(23) except that GFP was removed from the vector for compatibility with EKAREV. Doxycycline-inducible MEK^EE^ expression plasmid (pCW57-MEK^EE^) was constructed by subcloning constitutive MEK1 mutant, MEK^EE^ into pCW57 vector^(55)^.

### Mice

Transgenic mice expressing EKAR-EVnls were obtained from Laboratory Animal Resource Bank, National Institute of Biomedical Innovation, Health and Nutrition, Japan (56). Involucrin-tdTomato transgenic mice were obtained from Institute of Molecular Genetics of the ASCR, v. v. i., Czech Republic. All experimental procedures were carried out under the terms of a UK Home Office project license (PPL 70/8474) after local ethical review at King’s College London.

### Cell culture

Primary neonatal human keratinocytes (NHKs, strain km) were used in all experiments at passage 6-8. All cell stocks were routinely tested for mycoplasma contamination and were negative. The cells are not subjected to STR profiling because they are not an established cell line. Cells were cultured on a mitotically inactivated feeder layer of J2-3T3 cells in FAD medium (one part Ham’s F12, three parts Dulbecco’s modified Eagle’s medium, 1.8 × 10^-4^ M adenine), supplemented with 10% fetal calf serum (FCS) and a cocktail of 0.5 µg/ml of hydrocortisone, 5 µg/ml insulin, 10^-10^ M cholera enterotoxin and 10 ng/ml epidermal growth factor (HICE cocktail) (complete FAD medium)(17). For Ca^2+^ depleted keratinocyte culture medium, FCS was chelated with Chelex-100 resin (BioRad) and added to Ca^2+^ free FAD medium. J2-3T3 cells were cultured in high-glucose DMEM (Sigma-Aldrich) supplemented with 10% (v/v) adult BS (Life Technologies). HEK293T cells were cultured in high-glucose DMEM (Sigma-Aldrich) supplemented with 10% FBS (fetal bovine serum).

In some experiments NHKs were plated on collagen-coated flasks (pre-coated with rat tail collagen type I (Corning) at 20 µg/ml for 3 hours) and cultured in feeder-free conditions in keratinocyte serum-free medium (KSFM) containing 30 µg/ml BPE (bovine pituitary extract) and 0.2 ng/ml EGF (Gibco). KSFM-cultured cells were stimulated to differentiate (Fig.1 *F* and *G*) by exchanging the medium with high Ca^2+^ KSFM (1.2 mM) or complete FAD medium (containing 10% FCS).

Patterned PDMS substrates were pre-coated with rat tail collagen type I (Corning) at 20 µg/ml for 3 hours. Cells were seeded in complete FAD medium at a density of 75,000 cells cm^-1^ for 45 minutes at 37°C. Substrates were rinsed gently once with FAD medium to remove non-adherent cells and transferred to 6 cm dishes containing inactivated J2-3T3 cells seeded at a density of 20,000 cells/cm^2^.

EGF, TPA, and PD0525901 were added to the medium 6 hours prior to live imaging at final concentrations of 1 µg/ml and 1 µM, respectively.

### Patterned PDMS substrates

Patterned PDMS substrates were generated as previously described with circle diameter (*d*) 150 µm, center-to-center distance (λ) 300 µm, and UV light exposure time 20s(39).

### Lentiviral infection

All the transgenes were expressed in primary human keratinocytes by lentiviral transduction. Replication-defective and self-inactivating lentiviral vector (pCSII vector for EKAREV-nls and EKAREV-nes, pLenti vector for Involucrin-mCherry, pCW57 vector for doxycyclin-inducible DUSP expression) was co-transfected with packaging plasmid (pCAG-HIVgp) and VSV-G/Rev-expressing plasmid (pCMV-VSVG-RSV-Rev) into HEK293T cells (Clontech). Cells expressing Involucrin-mCherry were selected by 2 µg/ml Puromycin treatment.

### siRNA transfection

For knockdown of β1-integrin, DUSP6, and DUSP10, SMART pool ON-TARGET plus siRNAs (Ambion/ GE Healthcare) were used. The siRNAs were a mix of four sets of RNAi oligos.

For siRNA-mediated gene silencing, keratinocytes were cultured in KSFM containing 30 µg/ml BPE (bovine pituitary extract) and 0.2 ng/ml EGF (Gibco) for 2 days. Keratinocytes were transfected using INTERFERin (Polyplus transfections) with the final siRNA concentration of 30 nM and 4 µl INTERFERin reagent.

### In Situ Hybridization

RNAscope Fluorescent Multiplex Assay (Advanced Cells Diagnostics, USA) was used for in situ hybridization of human MAPK1, MAPK3, DUSP6, and DUSP10. NHKs were cultured in 24-well multiple well plates with feeder layer to allow colony formation. Following removal of the feeder layer, NHKs were fixed by 4% paraformaldehyde and subjected to protocols provided by ACD. For multiplex detection, samples were hybridized with probes against MAPK1and MAPK3 (C1) together with probes against DUSP6 or DUSP10 (C2).

### Microscopic detection of In Situ Hybridization signals

Images were acquired on a Nikon A1R laser scanning confocal microscope with GaAsp detectors using a 40x CFI Plan Apo Lambda 0.95 NA objective (Nikon) and NIS-Elements (Nikon). Nuclear signal was excited by a 405 nm laser and detected by 450/50 nm emission filter. MAPK1 and MAPK3 signals were excited by 638 nm laser and detected by 655/25 nm emission filter. DUSP6 or DUSP10 signal was excited by a 560 nm laser and detected by a 595/50 nm emission filter.

### Doxycycline-inducible overexpression

Doxycycline-inducible gene expression constructs (pCW57-DUSP6, pCW57-DUSP10, pCW57-MEK^EE^) were lentivirally co-transduced into NHKs with EKAREV or Involucrin-mCherry.1 µg/ml Doxycycline was added to the medium to induce DUSP expression. Live imaging was performed from 6 hours after doxycycline treatment.

### Live imaging of human keratinocytes

2D culture images were acquired on a Nikon A1R laser scanning confocal microscope with GaAsp detectors using a 20x Plan Apo VC 0.75 NA objective (Nikon) and NIS-Elements (Nikon). Live cells were imaged in a temperature-controlled chamber (37°C) at 5% CO_2_. For nuclear staining, Hoechst 33342 was added to the culture medium 30 min prior to imaging at a final concentration of 5 µg/ml. Images were acquired every 5 min for up to 24 hours. For FRET imaging, the EKAREV biosensor was excited by a 445 nm laser, and 482/35 nm and 525/50 nm emission filters were used to acquire CFP and FRET images, respectively. Involucrin-mCherry was excited by a 560 nm laser and detected by a 595/50 nm emission filter. The Hoechst 33342 signal was excited by a 405 nm laser and detected by a 450/50 nm emission filter.

Images for cells on PDMS substrates were acquired on a Nikon A1R confocal/multiphoton laser scanning microscope 25x Apo LWD 1.1 NA objective. Mineral oil was used to cover the surface of the medium and prevent evaporation. Live cells were imaged in a temperature-controlled chamber (37°C) and pH was maintained with 15 mM HEPES buffer. Images were acquired every 15 min.

### Single cell proliferation assay

Mitotically inactivated J2-3T2 feeder cells were plated on collagen-coated 384 well glass bottom plates in complete FAD medium at the density of 20,000 cells/cm^2^. Single NHKs were plated onto each well. 6 hours after plating, only wells that accommodate single cell were subjected to live imaging to measure ERK pulse levels for 6 hours. Cells were incubated for another 48 hours and the same wells were revisited by microscope to observe cell numbers.

### In vivo imaging of mouse epidermis

Methods for live imaging of mouse epidermis were previously described (11). Briefly, the ear skin of anaesthetized mice was depilated and stabilized between a cover glass and thermal conductive silicon gum sheet. In vivo imaging of tail epidermis was performed similarly. Depilated tail was flanked with two silicon gum sheets and a cover glass was placed on the top. In vivo live imaging was performed by ZEISS 7MP multi-photon microscope, equipped with W Plan-Apochromat 20x/1.0 DIC VIS-IR M27 75mm water-immersion objective lens and Coherent Chameleon Ti:Sapphire laser. EKAR-EVnls signal was detected by BP 500-550 and BP 575-610 for CFP and FRET, respectively.

### FRET analysis

Single cell ERK activity was measured by ratiometry of CFP and FRET signals (FRET/CFP) since EKAREV functions by intramolecular FRET and the molecular number of the two fluorescence proteins are considered to be equal. Each signal level was measured by mean pixel intensity in individual cell areas.

### Automated cell tracking

Tracking was performed by script-based operation of a FIJI plugin, Trackmate (http://imagej.net/TrackMate). FRET channel images were used for object detection and linking with manually optimized parameters. Identified object regions were redirected to corresponding CFP, FRET, and mCherry channel images to obtain mean intensities for each region. The whole data set of XY location, time, and mean intensities were exported to Excel software (Microsoft Corporation, Redmond, WA) or MATLAB 2018Ra software (Mathworks, Natick, MA) for further numerical analyses and data visualization.

### Quantification of In Situ Hybridization signal

The particle signals acquired by In Situ Hybridization were segmented by Trackmate. Individual cells were segmented by Watershed segmentation of NHK colony area based on nuclear positions. The whole data set of XY location, mean intensity of In Situ Hybridization signal and XY location of individual cells were exported to Excel software (Microsoft Corporation, Redmond, WA) or MATLAB 2018Ra software (Mathworks, Natick, MA) for further numerical analyses and data visualization.

### Semi-automated single-cell tracking of cells expressing EKAR-EVnes

Cells expressing EKAR-EVnes cultured on the patterned PDMS substrate (Fig. 4*D*) were tracked in a semi-automated manner with a custom-made program for FIJI/ImageJ. Cellular centre locations were tracked by eye based on the lack of nuclear signal of EKAR-EVnes. 12-pixel square regions were automatically created around each centre and Huang’s fuzzy thresholding was applied to obtain cytoplasmic regions expressing EKAR-EVnes. The mean CFP was FRET signals were obtained for each region and used for ERK activity (FRET/CFP).

### Segmentation of tip and base areas on the patterned PDMS substrate

Tip and base regions were demarcated by circles with a diameter of 200 μm centred at each tip (Fig. 4*D*, white dotted circles). ERK^high^ and ERK^low^ cells in base areas (troughs) were gated by the 1.2 value of FRET/CFP (Fig. 4 *E*-*G*).

### Instantaneous variance as a measure of population ERK pulse level

To quantify the level of ERK activity pulses of a population of cells at a given time point we measured the variance of ERK activity (FRET/CFP) at that time point among all cells (instantaneous variance). An increase in the instantaneous variance indicates a higher variability of ERK activity in the population at a specific time point. Pulses were observed to behave as stochastic events, as suggested by the exponential distribution of interpulse intervals (Fig. S1*D*). This, combined with the fact that the mean ERK activity did not change between conditions (Fig. 4*A*, Control/DUSP6) justifies the use of the instantaneous variance as a measure of the level of ERK activity pulses of a population at a specific time point. When the instantaneous variance remains unchanged between conditions (Fig. 4*B*, Control/DUSP10) we say that the level of ERK activity pulses are the same for both populations, regardless of changes in the mean level.

### Moving variance as a measure of ERK pulse level in a time window

In order to study the change in ERK activity pulses over time in individual cells, we analysed overlapping moving time windows of 50 minutes. Each time window was small enough for the mean ERK activity, within the window, to be considered fixed, but long enough to accommodate an ERK activity pulse (typical pulse ∼0.25 hr). For each window we computed the variance of ERK activity. The variance is a measure of dispersion of the measurements in the window, and quantifies the extent of the deviations of the signal from its mean value. These deviations can occur due to pulses, or noise (which is of much smaller amplitude than pulses). The variance captures both the amplitude and number of pulses in the time window, giving a quantitative measure of the pulsing level of ERK activity in the time window. The minimum value for the variance is zero, which corresponds to a signal without pulses or fluctuations; larger values of the variance indicate a higher level of pulsation.

This method allows a continuous assessment of the pulse level over time. Other methods, such as peak detection and pulse count, amount to a discrete measurement of pulses, which does not lend itself naturally to a detailed quantitative analysis of the temporal evolution of ERK activity pulses.

### Phase diagram

To construct the phase diagram of ERK activity variance and Involucrin mean level, cells co-expressing EKAREV-nls, and the Involucrin reporter, Involucrin-mCherry were considered. Time series obtained from automated cell tracking were then analysed by computing the variance of the ERK activity and the mean level of Involucrin on a moving window of 50 minutes. Only time series of more than 90 minutes were analysed.

The values of variance of ERK activity and mean Involucrin level were plotted for every time window, providing a trajectory in the plane spanned by ERK activity variance and mean Involucrin level. This was repeated for all cells.

The ERK activity variance vs Involucrin mean level plane was then divided into regular blocks. The trajectories that lay within each block were then averaged to obtain a mean direction for each block. This procedure resulted in the phase diagram of ERK activity variance vs mean Involucrin level. Every block in the phase diagram corresponds to a pair of ERK activity variance/Involucrin mean level values, while the arrow in the block indicates the mean direction to which these values changed in time.

The same procedure can be followed to construct other phase diagrams, such as mean ERK activity vs mean Involucrin level.

### Phase diagram normalization and transition probabilities

Each arrow in the phase diagram was decomposed into its x and y components. The x components were rescaled by the maximum value of the Involucrin mean level, i.e. x’=x/max(Inv. Mean level), while the y components were rescaled by the maximum value of the ERK activity variance, i.e. y’=y/max(ERK activity variance). This rescaling amounted to normalising both axes of the phase diagram to the range [0,1], and allowed the comparison between the x’ and y’ components of the arrows.

The transition probabilities between neighbouring blocks corresponded to r_x_=x’/(|x’|+|y’|) and r_y_=y’/(|x’|+|y’|), where |.| indicates the absolute value. Here, r_x_ accounts for the probability of transitioning to the neighbouring blocks of Involucrin mean level, while, r_y_ corresponds to the probability of transitioning to the neighbouring blocks of ERK activity variance. The sign of r_x,y_ indicates the direction of the transition, a minus (plus) sign signifies a transition towards decreasing (increasing) values of Involucrin/ERK activity variance. The probabilities are normalised, such that |r_x_|+ |r_y_ |=1.

### Cluster analysis

To analyse the spatial organisation of cells in ear and tail epidermis we measured the radial distribution function (RDF) g(*r*), which measure the deviations of the density of cells from that of a random distribution, as a function of the distance *r* from a cell of reference (57). To construct the RDF we constructed rings of radius *r* and width Δ*r* around every cell of interest, and counted total number N(*r*, Δ*r*) of cells that lie within the rings. We then constructed the reference number N_ref_(*r*, Δ*r*), which was computed by measuring N(*r*, Δ*r*) for a set of randomly distributed point in the field of interest. The field of interest was constructed by considering only regions of space where cells were observed in the experiment, as indicated by the black outline in Fig. 6 *G* and *M*, and S5 *C* and *D*. This was done to prevent artefacts due to boundary effects that might bias the clustering results. Finally, we constructed the RDF as the ratio g(r) = N(*r*, Δ*r*)/N_ref_(*r*, Δ*r*).

We constructed the null RDF g_null_(*r*)= N_null_(*r*, Δ*r*)/N_ref_(*r*, Δ*r*), against which the experimental measurements were be compared. In this case, both N_null_(*r*, Δ*r*) and N_ref_(*r*, Δ*r*) were measured for random distributions of points. In the null model, all particles are independent, randomly distributed points, hence the g_null_(*r*) is equal to unity. From 50 realisations of g_null_(*r*), we computed the 95% confidence intervals. When the experimental RDF lied above (below) the 95% c.i. the cells were considered clustered (dispersed) for that particular distance *r*, around cells.

The RDF was also used to study the clustering between two distinct populations, high and low pulsing cells. For this, we considered the cells of one group (high pulsing) as the reference cells around which the rings are constructed, while we counted the number of cells of the second group (low pulsing) that lie within the rings. In this case, when the experimental observations lie below (above) the 95% c.i. both cell populations are considered segregated (grouped).

### Statistics

Statistical analyses were performed using MS Excel or MATLAB R2018a (Mathworks) Software. We made use of the two-tailed Student’s t-test for unpaired data to quantify differences between experimental groups. Kolmogorov-Smirnov test was used to compare distributions. P values are indicated by: * 0.01<p<0.05, ** 0.01<p<0.001, *** p<0.001. n.s. = not significant.

## Acknowledgements

We thank Dr. Kazuhiro Aoki at National Institute for Basic Biology, Okazaki Institute for Integrative Bioscience, Japan for his advice on ERK pulse quantification. We thank Petr Kasparek, Institute of Molecular Genetics of CAS, for providing Involucrin-tdTomato mouse. The imaging was carried out in the Nikon Imaging Centre, King’s College London.

TH has received funding from the European Union’s Horizon 2020 research and innovation programme under the Marie Sklodowska-Curie grant agreement No 704587. IB was supported by CONICYT, Chile, Beca de Doctorado en el Extranjero No. 72160465, and the Centre for Doctoral Training on Theory and Simulation of Materials at Imperial College London, EPSRC (EP/L015579/1). FMW gratefully acknowledges financial support from the Medical Research Council (MR/PO18823/1), Wellcome Trust (206439/Z/17/Z) and BBSRC (BB/M007219/1).

---

**Fig. S1.**
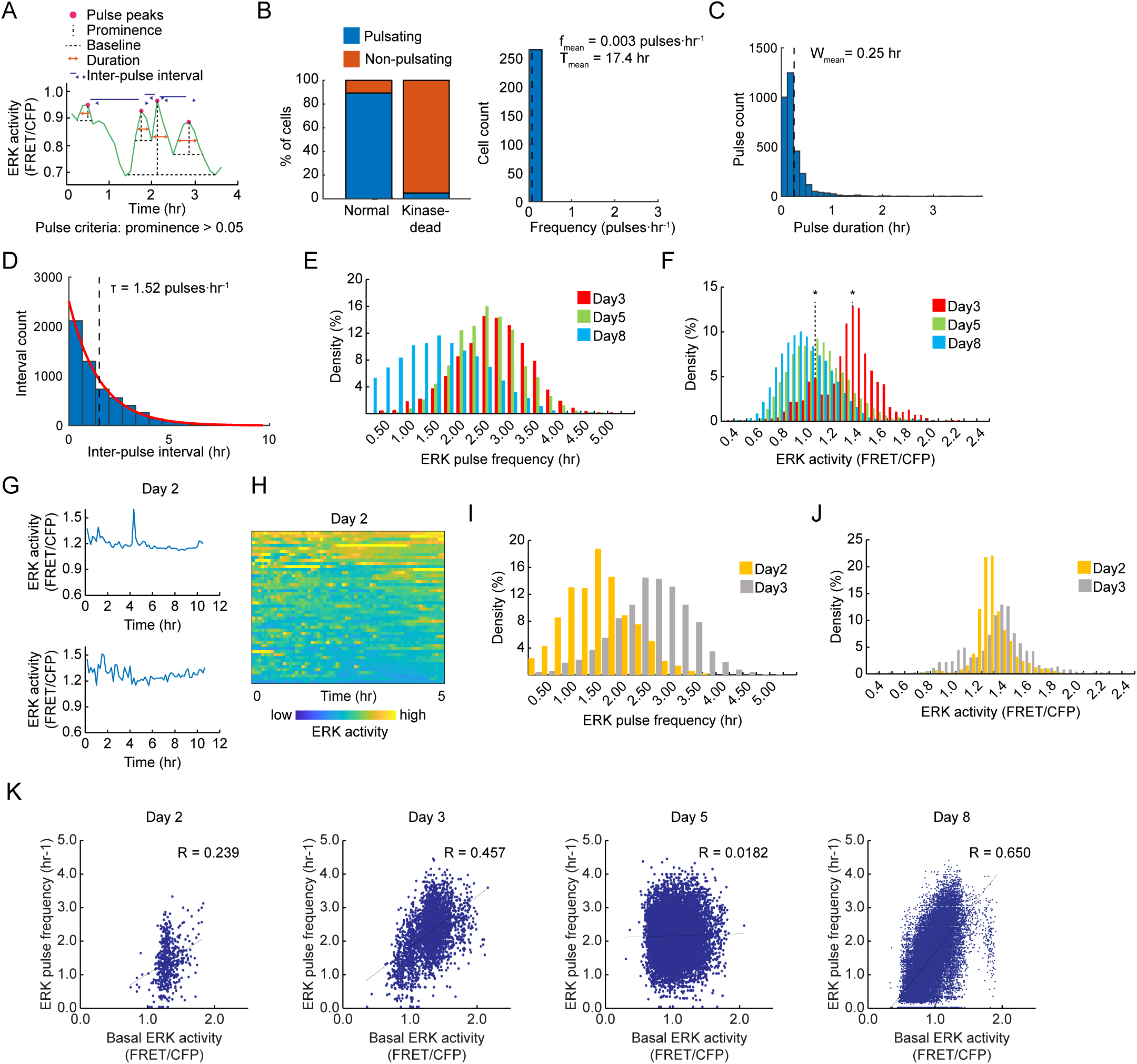
Quantitative analysis of ERK activation pulses. (*A*) Schematic of the ERK pulse detection and quantification. Pulses are detected as local peaks with prominence larger than 0.05 FRET/CFP value. Pulse duration was determined as the width of pulse at half the prominence of each pulse. Interpulse interval was characterised as the latency period between pulses. (*B*) Validation of the quantification methods with kinase-dead EKAR-EVnls biosensor (EKAREV-TA-nls), where FRET does not occur. Left, proportion of pulsatile cells in keratinocytes expressing normal EKAR-EVnls or EKAREV-TA-nls. Right, histogram of ERK pulse frequencies in pulsatile cells detected in keratinocytes expressing EKAREV-TA-nls. Data were obtained from human keratinocytes, cultured on feeder layers in complete FAD medium. (*C*) Histogram of pulse durations, indicating the mean pulse width. (*D*) Histogram of interpulse intervals fitted to an exponential decay curve, showing the value of the decay rate τ. (*E* and *F*) Histogram of ERK pulse frequency (*E*) and basal ERK activity (*F*) in keratinocytes on Day3, Day5 and Day8. Asterisks indicate the two peaks in the histogram of ERK activity on Day3 (red) (*n* = 3,323 cells for Day3, 11,527 cells for Day5, 37,320 cells for Day8 cells). (*G*) Representative time-series of ERK activity of NHKs on Day2. (*H*) Heat-map of ERK activity over time for 50 cells ordered by descending mean ERK activity (FRET/CFP) over time. Colours indicate ERK activity. (*I* and *J*) Histogram of ERK pulse frequency (*I*) and basal ERK activity (*J*) in keratinocytes on Day2 compared to Day3 NHKs. (*n* = 542 cells for Day2, 3,323 cells for Day3). (*K*) Dot plots and correlation analysis of basal ERK activity and ERK pulse frequency in NHKs on the indicated days. Lines indicate regression lines and values are Spearman’s rank correlation coefficient. (*n* = 542 cells for Day2, 3,323 cells for Day3, 11,527 cells for Day5, 37,320 cells for Day8 cells).

**Fig. S2.**
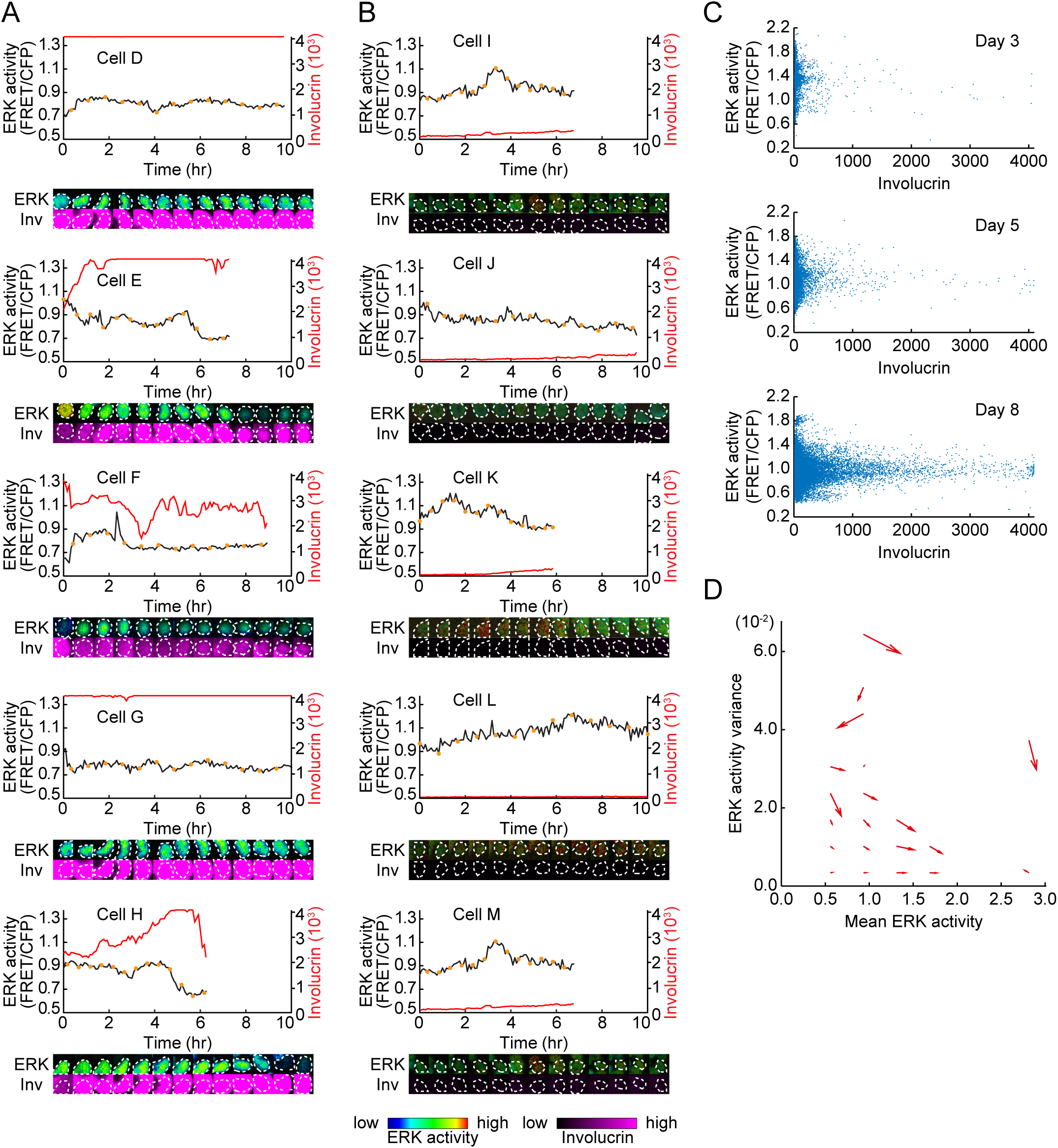
ERK activity profile in stably high and low Involucrin expression. (*A* and *B*) Representative time-series of ERK activity (black) and Involucrin-mCherry expression (red) of cells expressing stably high (*A*), or low (*B*) Involucrin. Images of the cells of ERK activity and Involucrin are shown for the time points indicated by the orange circles in each time-series. Images are shown by the indicated LUTs below. (*C*) Dot plot of mean ERK activity and Involucrin-mCherry expression in HNK cells on different days (*n* = 3,323 cells for Day3, 11,527 cells for Day5, 37,320 cells for Day8 cells). (*D*) Phase diagram of ERK activity variance and mean activity obtained from *n* = 3,323 cells. Arrows indicate the direction of transition for each compartment.

**Fig. S3.**
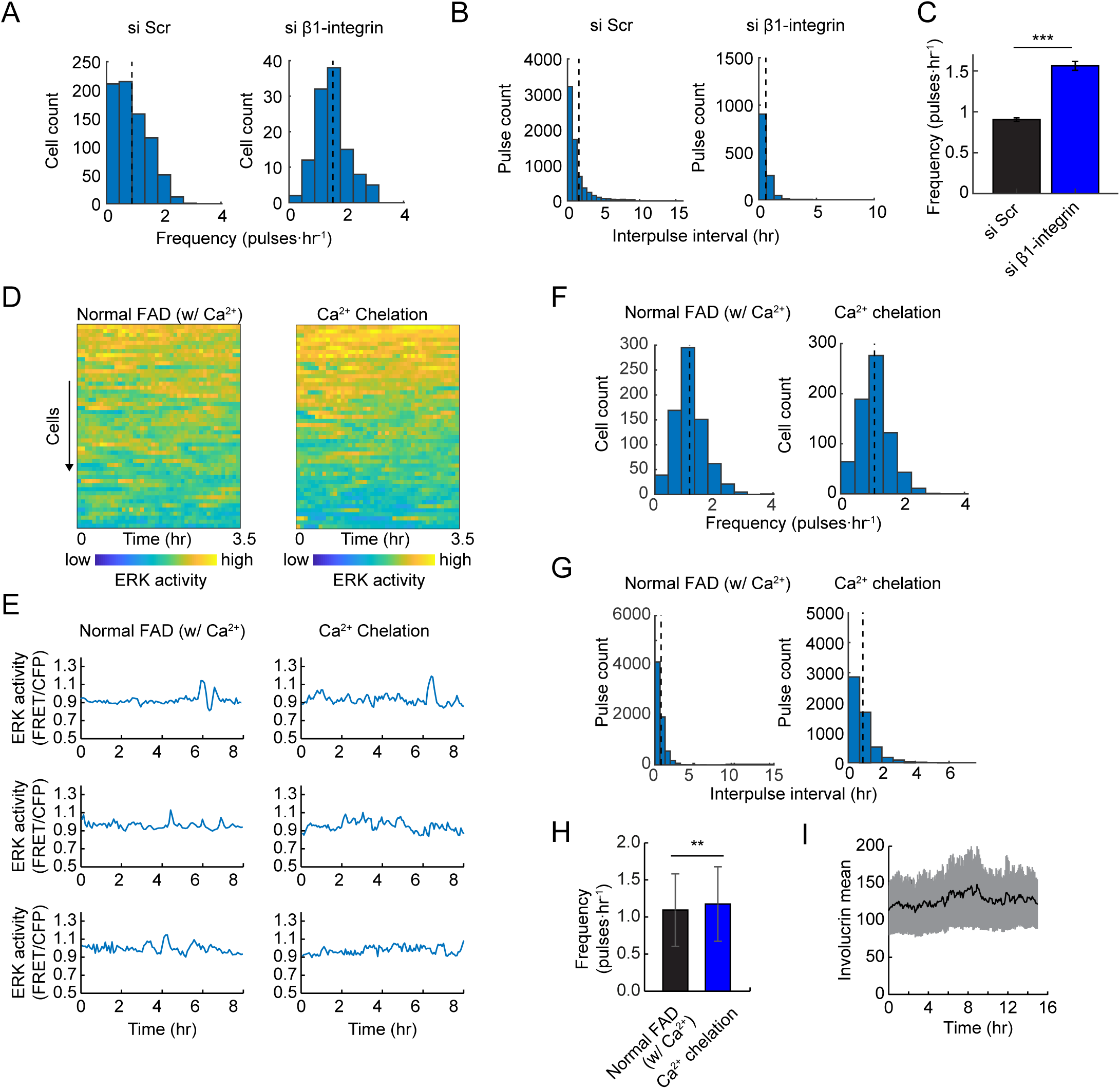
The effect of β1-integrin knockdown and Ca^2+^ chelation on ERK pulse patterns and differentiation. (*A* and *B*) Histograms of frequencies (*A*), and interpulse intervals (*B*), in cells treated with scrambled control siRNA (left) or β1-integrin-targeted siRNA (right). Black dotted lines: mean. (*n* = 764 cells for control siRNA, 112 cells for β1-integrin-targeted siRNA) (*C*) Frequency of ERK pulses. Data are shown by mean ± s.e.m (*n* = 764 cells for control siRNA, 112 cells for β1-integrin-targeted siRNA, Kolmogorov-Smirnov test; P=7.9 × 10^-20^). (*D*) Heat-map of ERK activity over time for 50 cells ordered by descending mean ERK activity (FRET/CFP) over time. Colours indicate ERK activity. (*E*) Representative time-series of ERK activity of cells treated with normal FAD medium (left) and Ca^2+^-chelated medium (right). (*F* and *G*) Histograms of frequencies (*F*), and interpulse intervals (*G*) in cells cultured with Ca^2+^-chelated medium. Black dotted lines: mean. (*n* = 757 cells for normal FAD treatment, 706 cells for Ca^2+^ chelated medium treatment). (*H*) Frequency of ERK pulses. Data are shown by mean ± s.e.m (*n* = 757 cells for normal FAD treatment, 706 cells for Ca^2+^ chelated medium treatment, Kolmogorov-Smirnov test; P=0.0245). (*I*) Mean Involucrin reporter expression over time. Data are shown as mean ± s.e.m. Statistical significance was examined by Kolmogorov-Smirnov test; P values are indicated by ** P<0.01, *** P<0.001.

**Fig. S4.**
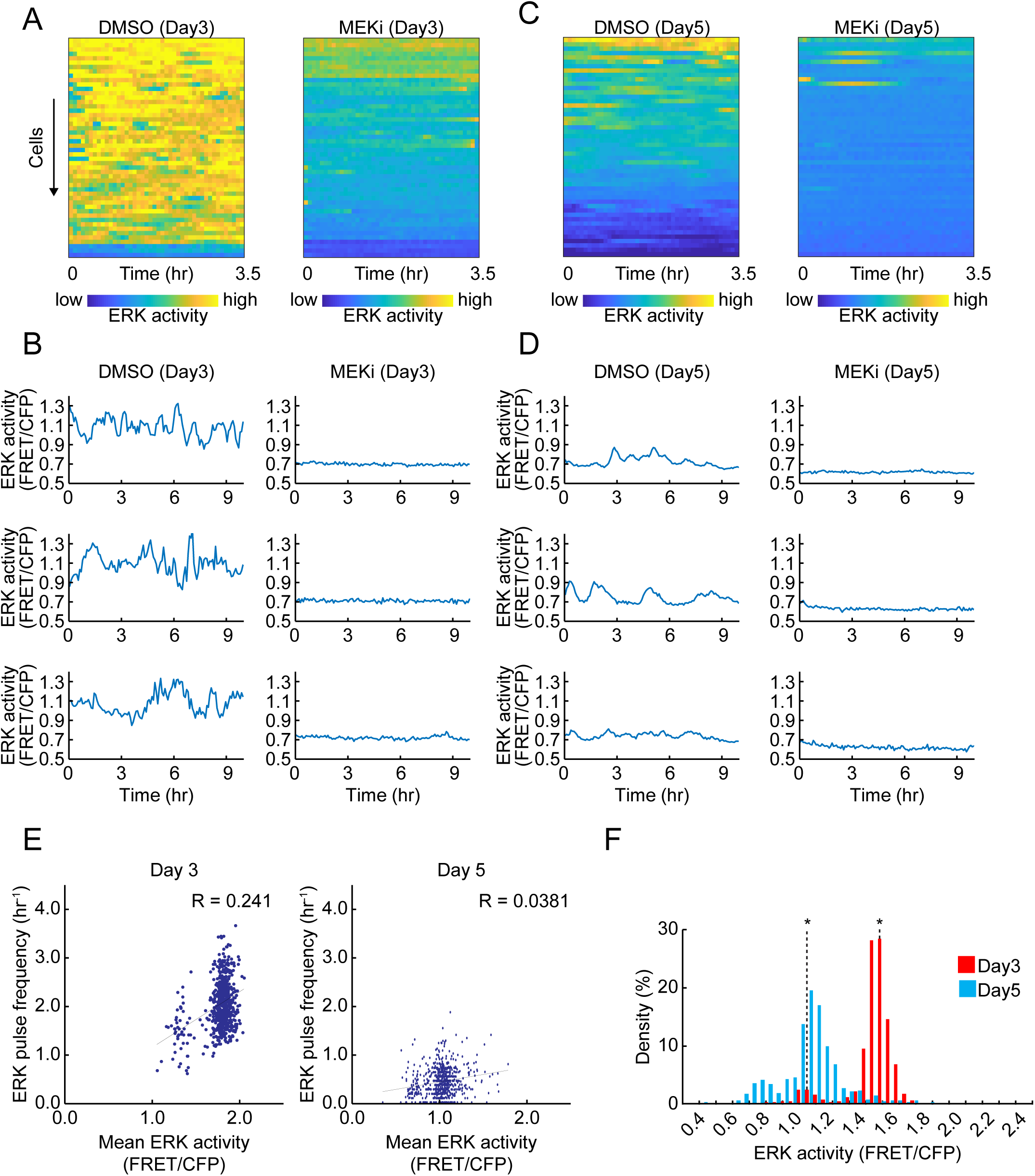
The effect of MEK inhibitor on ERK pulse patterns on different stem cell stages. (*A* and *C*) Heat-map of ERK activity over time for 50 cells ordered by descending mean ERK activity (FRET/CFP) over time. Colours indicate ERK activity. (*B* and *D*), Representative time-series of ERK activity of cells treated with DMSO or 1 μM MEK inhibitor, PD0325901 on day3 or day5 after plating. (*E*) Dot plots and correlation analysis of basal ERK activity and ERK pulse frequency in DMSO-treated NHKs on day3 (left) and day5 (right) after plating. Lines indicate regression lines and values are Spearman’s rank correlation coefficient. (*n* = 644 cells for Day3 cells, 675 cells for Day5 cells). (*F*) Histogram of ERK pulse frequency in NHKs treated with DMSO on day3 (left) and day5 (right) after plating. (*n* = 644 cells for Day3 cells, 675 cells for Day5 cells).

**Fig. S5.**
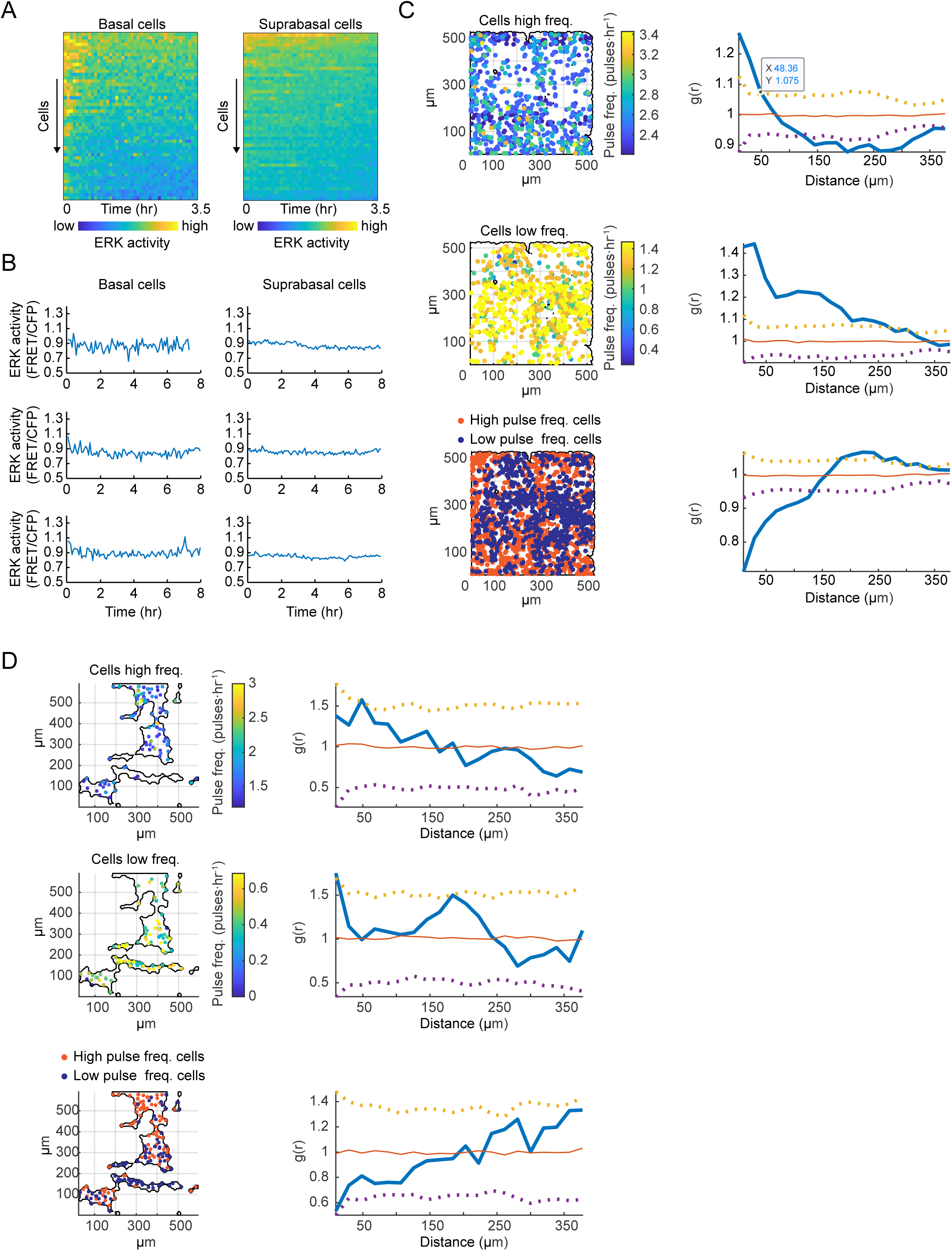
Clustering of cells with high or low ERK pulse frequency in mouse epidermis. (*A*) Heat-map of ERK activity over time for 50 cells ordered by descending mean ERK activity (FRET/CFP) over time in the basal (left) or suprabasal (right) layer of mouse epidermis. Colours indicate ERK activity. (*B*) Representative time-series of ERK activity in basal (left) or suprabasal (right) layer cells. (*C*) Clustering analysis of cells in the ear skin basal layer for the 700 most pulsatile (top) and 700 least pulsatile cells (middle) show the clustering between the most and least pulsatile subpopulations (bottom). The right panels show the radial distribution function g(r) (solid, blue line), null randomized (red lines), and the 95% c.i. (dotted lines). Values above or below the 95% c.i. indicate a significant clustering or dispersion of cells at the corresponding scale (distance), respectively. (*D*) The same analysis as (*C*), for the tail skin, where the 130 most and 130 least pulsatile cells were considered. The black boundaries in the left panels of (*C*), and (*D*), enclose the area used to compute the null lines.

**Movie S1**. ERK activity pulses in cultured human keratinocytes. Human keratinocytes expressing EKAR-EVnls cultured on feeder cell layer for 5 days. Colours indicate ERK activity. Image size: 624 μm × 624 μm.

**Movie S2**. Time-lapse movie of human keratinocytes on the patterned PDMS substrate. Human keratinocytes expressing EKAR-EVnes cultured on the patterned PDMS substrate for 48 hours. Colours indicate ERK activity. Image size: 507 μm × 507 μm.

**Movie S3**. Time-lapse movie of the basal layer of mouse tail epidermis. Mouse tail epidermis expressing EKAR-EVnls FRET biosensor. Colours indicate ERK activity. Image size: 332 μm × 332 μm.

